# T cell engagers convert acute tumor killing into endogenous vaccination

**DOI:** 10.64898/2026.10.06.756695

**Authors:** Mathilde M Lecuit, Béatrice Corre, Zacarias Garcia, Fabrice Lemaître, Amaury Vaysse, Ruby Alonso, Christel Goudot, Baptiste Arnaud, Lea C Feldmann, Carolina Moraes Cabé, Valérie Seffer, Pierre Guermonprez, Capucine L Grandjean, Marion V Guérin, Philippe Bousso

## Abstract

T cell engagers (TCEs) redirect T cells to kill tumors by cross-linking CD3ε to a tumor antigen, independently of T cell receptor specificity. Some patients achieve remissions that persist long after drug clearance, but whether TCEs can couple acute cytotoxicity to the generation of lasting adaptive immunity remains unknown. Here, using an immunocompetent mouse model for clinical-grade human TCEs, we show that TCE-mediated killing initiates an endogenous vaccination cascade. Intravital two-photon imaging reveals a perforin-dependent cytotoxic wave that peaks within 24 hours, then subsides despite massive T cell accumulation. Single-cell transcriptomics and adoptive transfers demonstrate that memory CD8⁺ T cells, the principal effector reservoir in mouse and human, transit through a progenitor-like state before differentiating into cytotoxic effectors with recall competence. Crucially, tumor lysis fuels cross-priming of tumor-specific CD8⁺ T cells by XCR1⁺ type 1 conventional dendritic cells, a cDC1-dependent step essential for durable tumor control and protective memory upon rechallenge.

## INTRODUCTION

The treatment of B cell malignancies has been transformed over the last decade by immunotherapies that redirect T cell cytotoxicity against tumors, by engineering antigen recognition independently of the endogenous TCR-peptide-MHC (pMHC) interaction^1–3^. Chimeric antigen receptor (CAR) T cells and bispecific T cell engagers (TCEs) embody this principle through distinct strategies: CARs graft synthetic antigen receptors onto autologous T cells, while TCEs are bispecific antibodies that simultaneously bind a tumor-associated antigen and CD3ε, forcing the formation of a cytotoxic synapse between tumor cells and the patient’s unmodified T cells^1,3,4^. CAR T cells have demonstrated remarkable efficacy in B cell malignancies but face significant constraints, including the time, cost, and complexity of patient-specific manufacturing, as well as the requirement for a lymphodepleting conditioning^5,6^. TCEs circumvent many of these barriers: as off-the-shelf molecules, they can be administered immediately to patients without prior cell collection, *ex vivo* engineering, or lymphodepletion^2,3^. This practical advantage, combined with the ability to engage polyclonal endogenous T cells *in situ*, has fueled substantial clinical interest^2,4^.

Since the first demonstration of clinical activity of a CD19-targeting Bispecific T-cell Engager (blinatumomab BiTE®) in hematological malignancies^7^, the field has expanded rapidly. Building on preclinical work demonstrating the potential of anti-CD20/CD3 bispecific antibodies^8^, several TCEs targeting CD20, CD19, or BCMA have now received clinical approval for relapsed or refractory B cell lymphoma and multiple myeloma. They have achieved remarkable response rates, including complete remissions that persist well beyond the expected duration of pharmacological activity^9–14^. Despite these significant clinical advances, the mechanism by which polyclonal, TCR-independent tumor lysis could lead to durable tumor control remains unresolved. In particular, whether TCEs can induce immunogenic cell death and thereby foster tumor-specific adaptive immunity remains a major open question.

Our mechanistic understanding of TCE activity derives almost entirely from *in vitro* studies, a setting in which TCEs promote polyclonal T cell activation, serial killing through perforin-granzyme release, and cytokine production^1,18–22^. More recently, single-cell transcriptomic analyses of patient samples have begun to characterize T cell states associated with TCE treatment^23–25^, and outline the prognostic value of various T cell subsets upon TCE initiation; the presence of pre-existing exhausted cells has been shown to be detrimental, while the maintenance of less-differentiated cytotoxic cells appears to be beneficial^14,24,26^. *In vivo*, several studies in immunocompetent models have reported rechallenge protection and memory T cell enrichment after bispecific antibody treatment^15,16^, including evidence of epitope spreading against antigens not targeted by the engager^15^. Yet the mechanism linking TCE-mediated killing to lasting immunity, and in particular whether dendritic cell-mediated cross-priming^17^ is involved, has not been addressed. More broadly, the *in vivo* dynamics of TCE-mediated killing, including the kinetics of T cell–tumor contact formation and the identity of T cell subsets that generate cytotoxic effectors, remain largely unexplored. This knowledge gap stems largely from the lack of suitable preclinical models: humanized mice engrafted with human peripheral blood mononuclear cells develop graft-versus-host disease and lack a complete immune microenvironment^27^, while existing strategies to humanize the CD3 locus in syngeneic models do not fully recapitulate the human TCE–T cell interaction^28,29^.

Here we develop an immunocompetent mouse model that overcomes these limitations and dissect how TCE drives tumor killing *in vivo*, from the early induction of a killing wave to the profound reprogramming of the T cell pool and, ultimately, the establishment of protective immunological memory.

## RESULTS

### An immunocompetent mouse model to study human T cell engagers *in vivo*

To study the mechanisms underlying the therapeutic effects mediated by human TCEs *in vivo*, we sought to develop a versatile and immunocompetent preclinical model in which murine T cells express the molecular target of human TCEs, the human CD3ε molecule (hCD3ε). We generated bone marrow chimeras by transducing donor hematopoietic stem cells with a retroviral vector encoding hCD3ε together with the transduction marker human CD34 (hCD34), and transplanted them into lethally irradiated recipients (referred to as WT^hCD3ε^ mice; **Figure 1A**).

**Figure 1.**
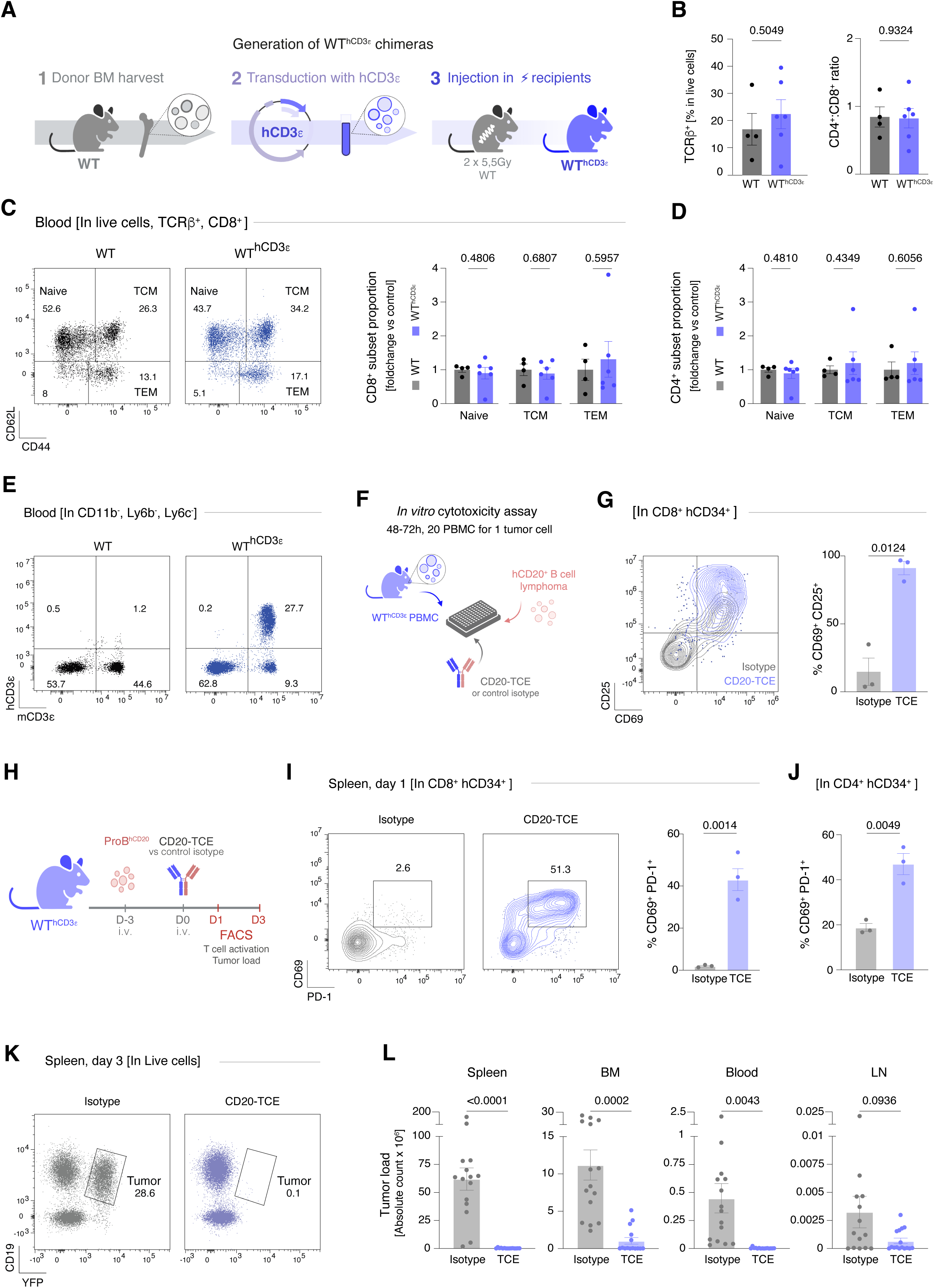
An immunocompetent mouse model for the exploration of human T cell engagers in vivo. **A**. Schematic of WT^hCD3ε^ chimera generation. Donor hematopoietic stem cells are harvested from the bone marrow, transduced with hCD3ε (and the transduction marker hCD34), then transplanted into lethally irradiated recipients. **B-D: Blood characterization of hCD3ε chimeras. B.** Frequency of TCRβ⁺ cells among live cells and CD4:CD8 ratio in the blood of WT (grey) and WT^hCD3ε^ (blue) chimeras (n = 4 per group). **C.** CD8⁺ T cell subset composition (naïve, TCM, TEM) in the blood (representative FACS plot and quantification as a foldchange to WT mean per subset). **D**. CD4⁺ T cell subset composition (naïve, TCM, TEM) in the blood (quantification as a foldchange to WT mean per subset). **E.** Expression of hCD3ε across CD11b^-^, Ly6b^-^, Ly6^-^ cells in the blood, demonstrating mCD3ε-cell-restricted surface expression. **F-G: *In vitro* validation. F.** Schematic of *in vitro* validation: PBMCs are harvested from the blood of WT^hCD3ε^ and co-cultured with hCD20⁺ murine B tumor cells in the presence of TCE or isotype control. **G.** Expression of activation markers CD69 and CD25 on hCD34^+^ CD8^+^ T cells (surrogate for hCD3ε^+^ CD8^+^ T cells) after 48-72 h co-culture with hCD20^⁺^ Pro-B, upon TCE or isotype control treatment (representative FACS plot and quantification). Each dot represents the mean of 3 technical replicates (wells) from one independent experiment. Bars indicate the mean ± SEM of n=3 independent experiments. **H-L: *In vivo* validation**. **H.** *In vivo* experimental design: WT^hCD3ε^ mice are injected intravenously with hCD20⁺ Pro-B cells on day -3 and treated with a single intravenous dose of 1mg.kg^-1^TCE or isotype control on day 0; FACS analysis is performed at days 1 and 3. **I.** CD8^+^ T cell activation on D1, assessed by CD69 and PD-1 co-expression on hCD34^+^ CD8⁺ T cells in isotype-versus TCE-treated mice (representative FACS plot and quantification). Each dot represents one mouse, from one experiment. **J**. CD4^+^ T cell activation on D1, assessed by CD69 and PD-1 co-expression on hCD34^+^ CD4⁺ T cells in isotype-versus TCE-treated mice (quantification). Each dot represents one mouse, from one experiment. **K.** Representative FACS plot of tumor load evaluation in the spleen on day 3 of treatment. Tumor is gated in live cells as YFP^+^ CD19^+^ cells. **L**. Quantification of absolute tumor cell count across spleen, bone marrow, blood and lymph nodes, on day 3 from treatment (TCE or isotype control). Each dot represents one mouse (n=15 isotypes, n=15 TCE). Data are pooled from 4 independent experiments. P values by two-tailed unpaired t test with Welch’s correction.

We first verified that this approach preserved normal T cell development and homeostasis. Eight to twelve weeks post-reconstitution, WT^hCD3ε^ chimeras exhibited thymic development resembling age-matched WT mice, with similar proportions of double-negative (DN), double-positive (DP), and single-positive (SP) subsets (**Extended Data Figure 1A**). In the periphery, T cell proportions, CD4⁺: CD8⁺ ratios, and the distribution of naïve, central memory (T_CM_), and effector memory (T_EM_) subsets were comparable to WT controls across blood (**Figure 1B–D**), spleen, and lymph nodes (**Extended Data Figure 1B–C**). Regulatory T cells were present in similar proportions (**Extended Data Figure 1D–E**). As expected, hCD3ε surface expression was restricted to cells expressing endogenous mCD3ε, as this process requires its integration within the CD3/TCR complex^30^ (**Figure 1E**). Most T cells robustly expressed hCD3ε, as assessed by hCD34 reporter expression (**Extended Data Figure 1F**). Thus, WT^hCD3ε^ mice maintain a physiological T cell compartment while broadly expressing the human target of TCEs.

We next asked whether T cells from WT^hCD3ε^ mice were functionally responsive to human TCEs. We transduced two murine B lymphoma cell lines (Pro-B and Eμ-Myc) to express human CD20, and co-cultured them with PBMCs from WT^hCD3ε^ mice in the presence of distinct human CD20-targeting TCEs: a research-grade CD20-TCE^8^, clinical-grade glofitamab, or mosunetuzumab (**Figure 1F**). Upon TCE exposure, both CD8⁺ and CD4⁺ T cells upregulated activation markers including CD69, CD25, CD44, and PD-1 (**Figure 1G, Extended Data Figure 2A–D**), confirming functional engagement of murine T cells by human TCEs in our system.

Finally, we evaluated whether WT^hCD3ε^ chimeras supported TCE-mediated tumor control *in vivo*. Mice were injected intravenously with hCD20⁺ Pro-B tumor cells three days before treatment with a single intravenous injection of 1 mg.kg⁻¹ TCE or isotype control (**Figure 1H**). Control mice developed disseminated lymphoma that particularly invaded the bone marrow and spleen; in contrast, TCE induced rapid T cell activation (**Figure 1I–J**) and promoted widespread tumor clearance within three days across all invaded organs (**Figure 1K–L**).

Together, WT^hCD3ε^ chimeras represent an immunocompetent platform that preserves physiological T cell biology while enabling the mechanistic study of clinical-grade human TCEs *in vivo*.

### TCE triggers immediate interactions between tumors and T cells that subsequently develop LFA-1/perforin-dependent lethality

To dissect TCE-mediated anti-tumor activity *in vivo*, we first characterized the dynamics and outcomes of T cell-tumor interactions using intravital imaging. To visualize endogenous T cells, we introduced a fluorescent component into our chimeric mice: Lck-Cre × Rosa26-mTmG bone marrow cells were transduced with hCD3ε to obtain *Lck*GFP^hCD3ε^ chimeras, whose T cells expressed membrane GFP as well as hCD3ε (**Extended Data Figure 3A**). These animals were engrafted with hCD20⁺ malignant B cells expressing a FRET-based fluorescent reporter for apoptosis^31^, allowing the simultaneous visualization of T cell–tumor interactions and caspase-3-dependent apoptosis in tumor cells upon TCE administration. We focused on changes and interactions upon TCE injection in the bone marrow, a primary site of tumor invasion (**Figure 2A**).

**Figure 2.**
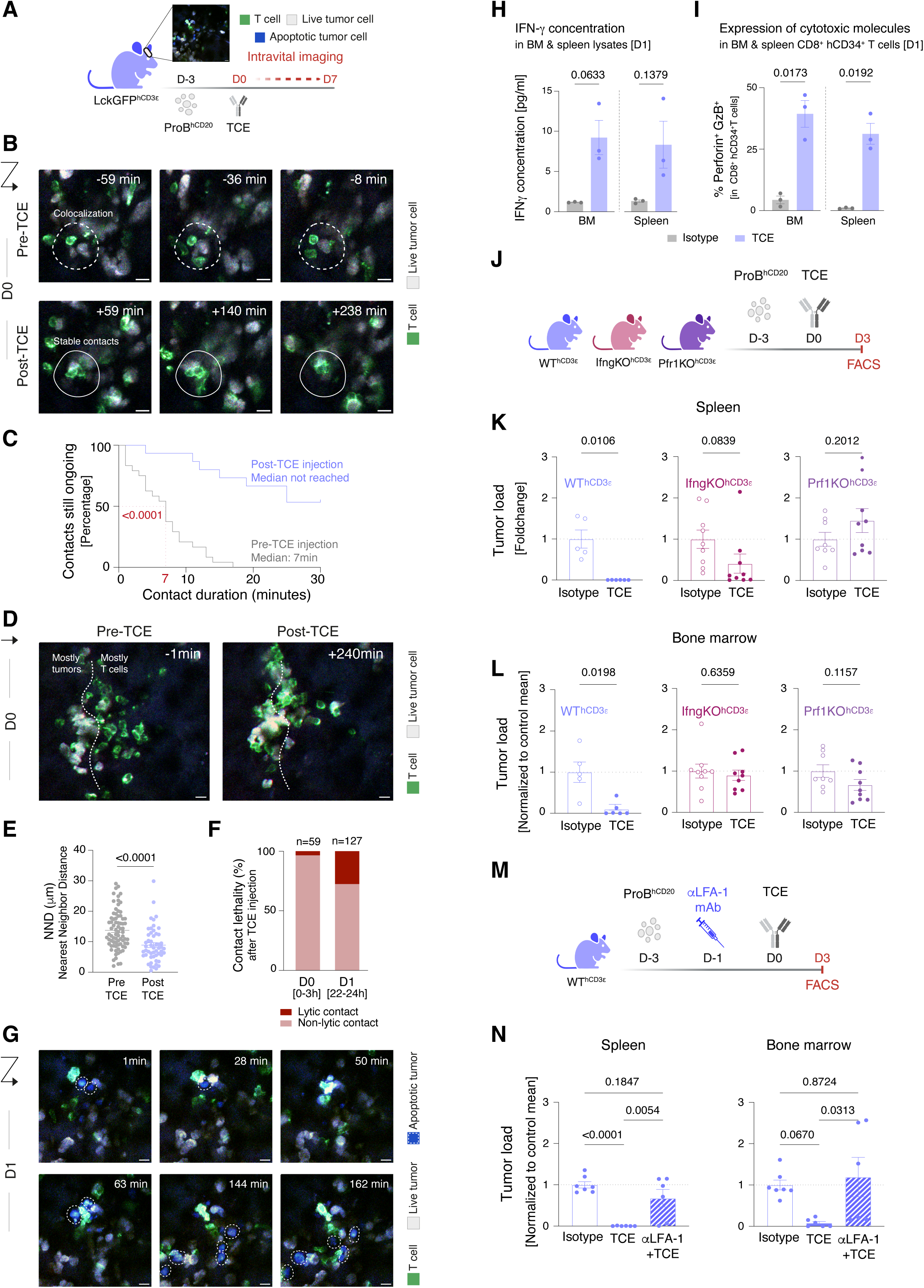
TCE promotes formation of LFA-1/perforin-dependent cytotoxic contacts with delayed killing competence. A-G: Intravital imaging in LckGFP^hCD3ε^ chimeras. **A.** Experimental design: LckGFP^hCD3ε^ chimeras bearing hCD20⁺ tumor cells expressing a FRET-based apoptosis reporter are imaged by intravital two-photon microscopy of the calvarium bone marrow. **B.** Representative intravital snapshots of one BM position before (pre-TCE) and after (post-TCE) injection. Green: T cells; gray/white: live tumor. Timestamps indicate time relative to film initiation (pre-TCE) or after treatment injection (post-TCE). **C.** Contact duration survival analysis: the percentage of T cell-tumor contacts still ongoing is plotted as a function of contact duration. Before TCE (grey), contacts are short-lived (median duration 7 minutes). After TCE (blue), contacts are markedly stabilized, with more than 50% persisting beyond the 30-minute observation window (P < 0.0001, log-rank test). **D.** Representative intravital snapshots of one BM position, pre- and post-TCE injection. Green: T cells; gray/white: live tumor. Dotted line: artificial limit between T cells and tumor cells before TCE administration. Timestamps indicate time relative to TCE injection. **E.** Nearest-neighbor distance (NND) between each tumor cell and the closest T cell, measured on the first and last frame of a time-lapse movie. Each dot is a tumor cell (n = 76 at the start, n = 52 at the end); data were pooled from 3 different positions. Bars show mean ± SEM. Unpaired two-tailed Welch’s t-test, p < 0.0001. **F.** Proportion of T cell–tumor contacts resulting in tumor lysis after TCE injection, on D0 (from 1 to 3 hours) versus D1 (22-24h). For each timepoint, the number of new apoptosis events was reported as a percentage of the total contact count during that timepoint (D0: 59 contacts, D1: 127 contacts). **G.** Representative intravital snapshots of one BM position on D1 after TCE injection. Top left to bottom right: presence of apoptotic tumor cells at the start of film acquisition (minute 1) and the accumulation of apoptotic events over 3 hours of filming. Green: T cells; gray/white: live tumor; blue: apoptotic tumor. Timestamps indicate time relative to film initiation. Scale bars, 10μm. **H-N: Exploration of molecules involved in TCE-mediated killing through genetic and pharmacological impairment of T cell functions. H.** Early increase in IFN-γ concentration measured by ELISA in lysates of BM & spleen (day 1, pg/ml). Each dot represents one mouse. Data from one experiment. P values by two-tailed unpaired t-test with Welch’s correction. **I.** Early expression of cytotoxic molecules by CD8^+^ T cells as represented by the percentage of double positive (perforin & granzyme B) CD8^+^ T cells among CD8^+^ hCD34^+^ T cells in the BM & the spleen, on day 1. Each dot represents one mouse. Data from one experiment. P values by two-tailed unpaired t-test with Welch’s correction. **J.** Schematic of genetic perturbation experiments: WT^hCD3ε^, *Prf1*KO^hCD3ε^ and *Ifng*KO^hCD3ε^ chimeras are treated with TCE or isotype control before evaluation of the tumor load on D3. **K-L:** Tumor load in spleen **(K)** and bone marrow **(L)** following TCE treatment in WT^hCD^^3^^ε^, *Prf1*KO^hCD^^3^^ε^ and *Ifng*KO^hCD^^3^^ε^ chimeras. P values by two-tailed unpaired t-test with Welch’s correction. **M.** Schematic of pharmacological perturbation experiment: WT^hCD3ε^ mice receive 1mg.kg^-1^ IV anti-LFA-1 (αLFA-1) blocking antibody one day before TCE administration. **N.** Tumor load in spleen and bone marrow in WT^hCD3ε^ mice treated with isotype, TCE, or TCE preceded by anti-LFA-1. Each dot represents one mouse. Data are pooled from 3 independent experiments. P values by one-way ANOVA with Tukey’s adjustment for multiple comparisons.

Before injection, T cells appeared motile and engaged in transient contacts with tumor cells, with a median duration of 7 minutes (**Figure 2B, top; Figure 2C; Extended Data Video 1**). Within minutes of TCE injection, T cell behavior shifted dramatically: contacts became stable and prolonged, with more than 50% of T cell-tumor conjugates persisting beyond the 30-minute analysis window (**Figure 2B, bottom; Figure 2C**). Many of these contacts were still ongoing hours after TCE injection (**Extended Data Video 2**). This resulted in the clustering of T cells within the tumor mass (**Figure 2D**) and a significant narrowing of the minimal distance between a given tumor cell and its closest T cell neighbor (**Figure 2E**). Despite the formation of these prolonged contacts, only ∼3% of contacts resulted in killing events within these first few hours. By contrast, lysis of tumor cells by T cells was markedly enhanced at 24 hours post-injection, at which point almost 30% of T cell contacts were lytic (**Extended Data Video 3, Figure 2F**). Of note, day 1 accumulation of apoptotic tumors in the bone marrow, as shown in **Figure 2G**, could also be detected in the spleen by flow cytometry (**Extended Data Figure 3B**). Thus, TCE drives a sequential reorganization of interactions between T cells and tumor cells: within minutes of injection at day 0, T cells form clusters within the tumor and establish long-lasting yet non-lytic contacts. Within 24 hours, T cells have acquired killing potential, prompting the widespread accumulation of apoptotic tumor cells.

We then sought to assess the contribution of different molecular pathways to the tumor killing observed upon TCE administration. In accordance with previous studies demonstrating the role of perforin/granzymes and cytokine secretion for TCE-mediated killing *in vitro*^18–22^, we found that TCE promoted degranulation of hCD3ε chimeric CD8^+^ T cells (**Extended Data Figure 3C)**. Moreover, we confirmed *in vivo* that within a day, TCE promoted IFN-γ production in the tumor microenvironment as well as accumulation of cytotoxic molecules in CD8^+^ T cells (**Figure 2H**, **I)**. We thus leveraged our chimeric model and generated *Prf1*KO^hCD3ε^ and *Ifng*KO^hCD3ε^ chimeras alongside WT^hCD3ε^ mice to evaluate their respective impact on early tumor killing (**Figure 2J**). While IFN-γ appeared to be mostly dispensable for tumor control in the spleen, perforin was required for TCE-induced killing in all invaded organs (**Figure 2K, L**). To evaluate if TCE was sufficient to generate and stabilize the cytotoxic synapse through which perforin and granzymes are delivered, we injected a blocking antibody against the synaptic integrin LFA-1 the day before TCE administration (**Figure 2M**). While blockade of LFA-1 did not impact the synthesis of cytotoxic molecules in T cells upon TCE treatment, it markedly impaired tumor killing (**Extended Data Figure 3D, Figure 2N**). Together, these data support a model where simultaneous binding of tumors and T cells by TCE is sufficient to trigger cytotoxicity in T cells, yet integrins are required for the execution phase: stabilization of the immune synapse by LFA-1 allows the release of granzymes and perforin to drive early tumor killing *in vivo*.

### Early killing wave is followed by unresponsiveness to TCE despite T cell accumulation

Whether TCE-induced killing is sustained or transient remained unclear. To assess this, we took advantage of intravital imaging to quantify apoptosis at later time points in hCD20^+^ Pro-B-bearing mice. To account for the decrease in tumor load upon treatment, we normalized apoptosis to total tumor cell area within each frame. Detection of apoptotic cells peaked at day 1 before a progressive decline through days 2, 3, and 7 (**Figure 3A, B**). Strikingly, the waning of apoptosis was accompanied by the massive accumulation of T cells within the tumor, with maximal density at day 3 before a sharp decrease at day 7 (**Figure 3C**). Therefore, acute killing after TCE appears to be transient, preceding a wave of T cell infiltration into the tumor.

**Figure 3.**
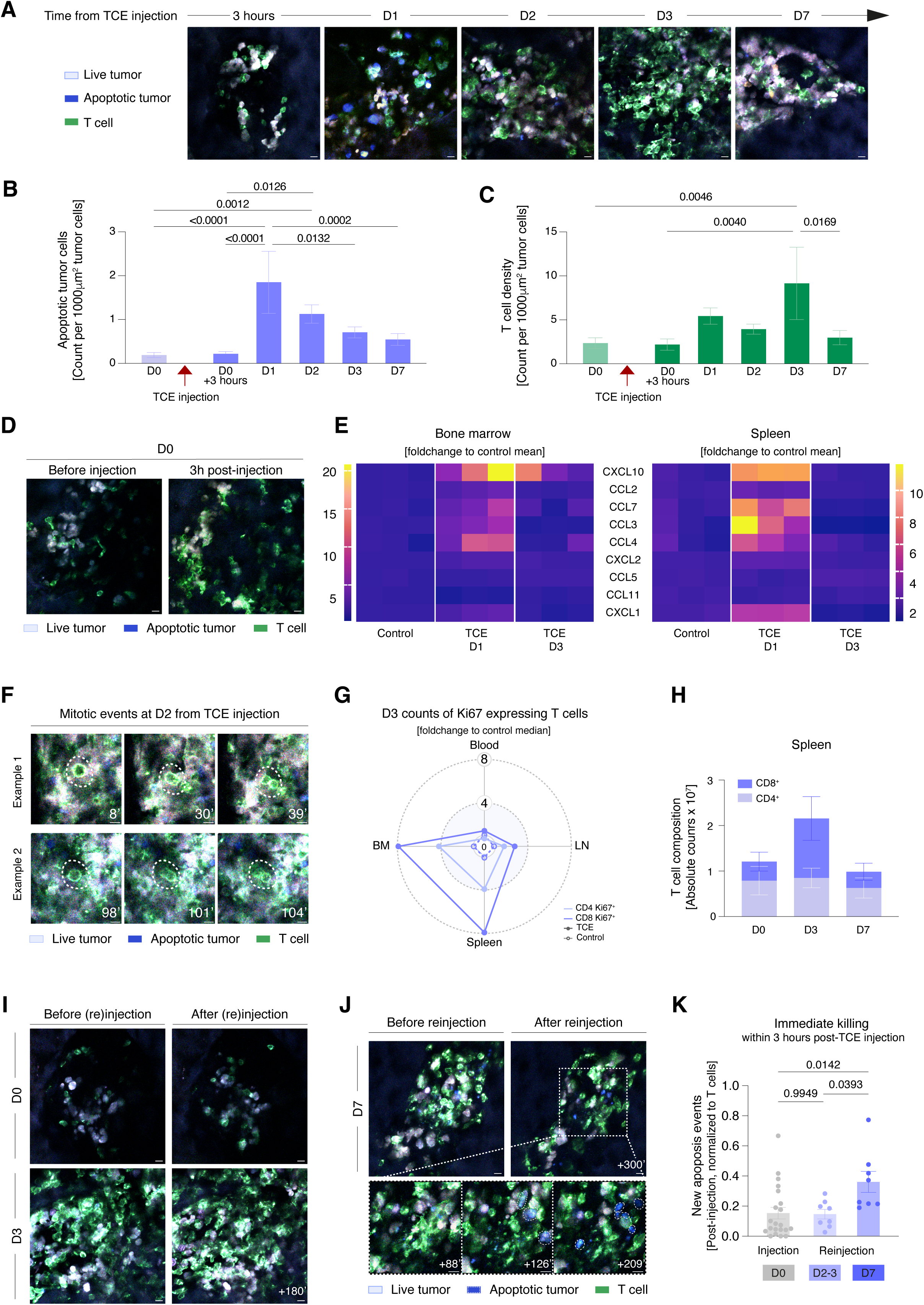
An early killing wave is followed by a refractory period and recall competence at day 7. **A-C. Intravital imaging revealing kinetics of tumor killing and T cell accumulation after TCE injection. A.** Representative intravital images of bone marrow at the indicated timepoints after TCE injection. Green, T cells; cyan/white, live tumor; blue, apoptotic tumor. Scale bars, 10μm. **B**. Quantification of apoptotic tumor cell density over time post-TCE, measured as the count of apoptotic cells normalized by total tumor density per frame to allow inter-timepoint comparison. **C**. T cell density within the tumor mass over time. **D-H: Exploration of the mechanisms driving T cell accumulation**. **D.** Example of one occurrence of T cell swarming within the tumor 3 hours after TCE injection. **E**. Heatmap of chemokine concentrations in spleen and bone marrow (fold-change relative to isotype controls) at day 1 and 3 post-TCE, measured via multiplex ELISA on tissue lysates. **F**. Representative intravital images capturing mitotic events at day 2. **G**. Ki67 expression in T cells across bone marrow, spleen, blood and lymph nodes (radar plot, fold-change relative to control median). For each organ, CD8^+^ T cells are dark blue, CD4^+^ T cells are light **blue, TCE-treated in full circles and lines, isotype control in empty circles and dotted line. H**. Absolute counts of CD8⁺ (dark blue) and CD4⁺ (light blue) T cells in the spleen on days 0, 3 and 7 post-TCE. **I-K: Temporal gating of killing boost upon TCE reinjection. I**. Representative intravital images showing the outcome of TCE injection on days 0 (T cell relocation) and 3 (no killing despite abundant T cells). **J**. Same as (*I*) but with re-injection on day 7 (immediate tumor lysis). Timestamps indicate time relative to injection (D0) or re-injection. (D3 & 7) Scale bars, 10μm. **K**. Quantification of apoptotic events per imaging field following TCE re-injection at the indicated timepoints. Each dot represents one position, pooled from 2-3 independent imaging sessions per timepoint. P values by one-way ANOVA with adjustment by Tukey’s multiple comparison test (**B**, **C**, **K**).

We next sought to dissect the mechanisms driving this increase in T cell density after TCE. Intravital imaging detected some events of T cell swarming as early as 3 hours post-injection (**Figure 3D**). We hypothesized that chemokine secretion could account for the rapid redistribution of T cells, as previously described with CAR-T cells^32^. To test this hypothesis, we performed multiplex ELISA profiling of the spleen and bone marrow (**Figure 3E, Extended Data Figure 4A, B**). In both compartments, TCE triggered a rapid chemokine burst at day 1, more particularly CXCL10, CCL3, and CCL4, previously implicated in T cell recruitment^32,33^. Accumulation was further sustained by local proliferation, as evidenced by intravital imaging, which detected mitotic events within the tumor at day 2 (**Figure 3F, Extended Data Video 4**). In addition, widespread upregulation of Ki67 was detected in T cells, particularly in the CD8^+^ subset (**Figure 3G**). This translated into a threefold expansion of CD8⁺ T cells at day 3 followed by contraction at day 7, while CD4⁺ T cell counts remained unchanged throughout (**Figure 3H**).

Because the accumulation of CD8^+^ T cells was concomitant with a decrease in tumor killing, we asked whether the cytotoxic deficit could be overcome by re-administration of TCE. Thus, we challenged mice with a second injection during intravital film acquisition. Strikingly, while T cell density was maximal on days 2 and 3 after the first injection, re-injection of TCE at this early timepoint did not induce tumor lysis by the surrounding T cells (**Figure 3I**). In contrast, administration of a second dose at day 7 triggered immediate tumor killing, suggesting that T cells had acquired enhanced cytotoxic competence during the refractory period (**Figure 3J, K, Extended Data Video 5**).

Together, these results show that TCE induces an early but transient killing wave, bolstered by a chemokine burst that contributes to an accumulation of T cells that appear to be temporarily unresponsive to TCE. Our data suggest the acquisition of recall cytotoxicity is not determined by CD8⁺ T cell counts but rather time-controlled, pointing to an active reprogramming of the CD8⁺ T cell compartment.

### The refractory period reflects active CD8^+^ T cell reprogramming toward recall-ready cytotoxic effectors

To characterize how TCE transforms the CD8⁺ T cell compartment to shape its lethality over time, we performed single-cell RNA sequencing of splenic cells harvested at days 0, 3 and 7 after TCE.

Single-cell RNA-seq recapitulated the transient expansion of CD8⁺ T cells at day 3 and revealed that TCE-induced cells were transcriptionally remarkably distinct from steady-state populations (**Figure 4A-B**).

**Figure 4.**
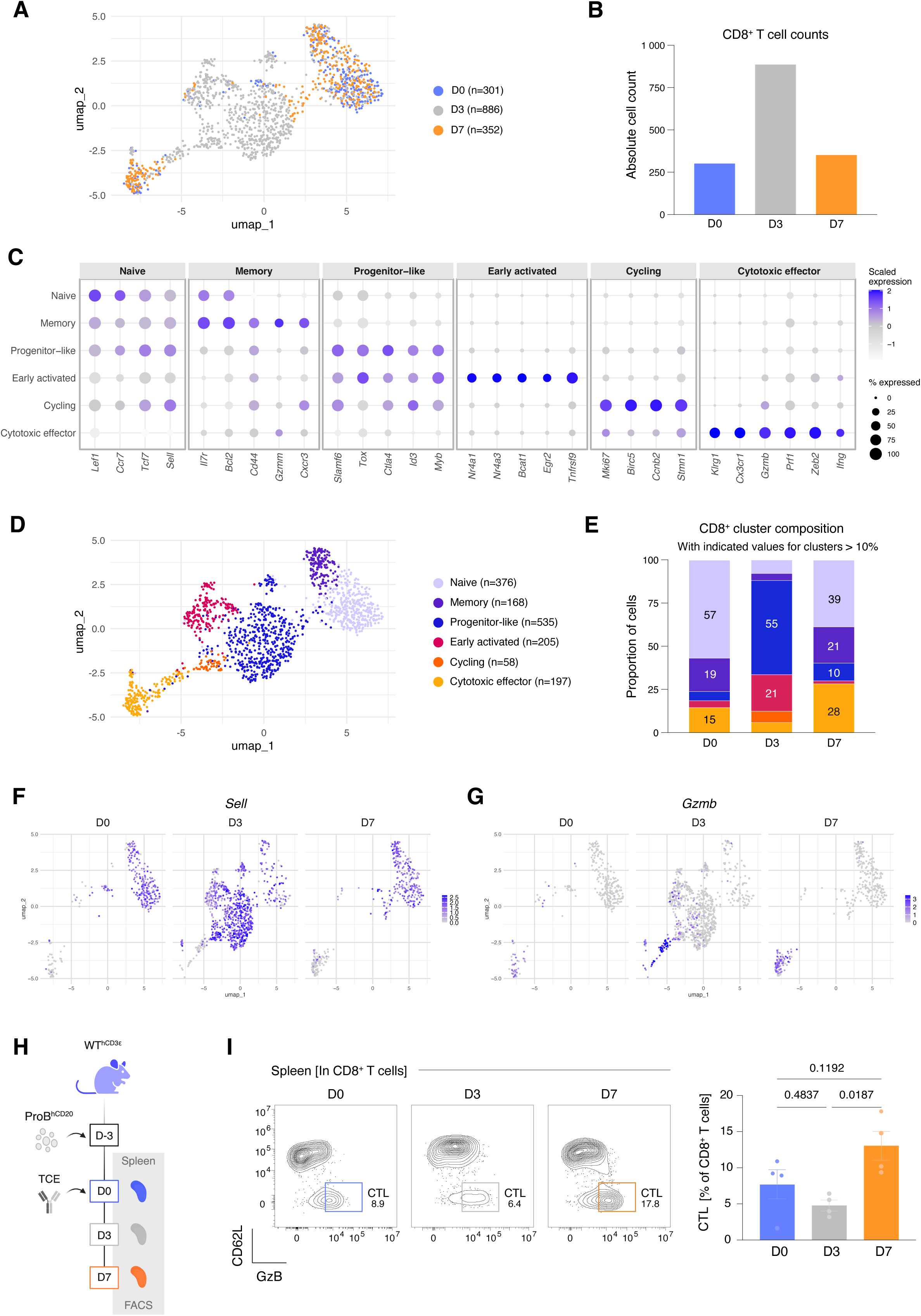
TCE drives CD8⁺ T cell reprogramming through a progenitor-like intermediate. A-G: Analysis of CD8^+^ reprogramming through single-cell RNA-sequencing. **A**. UMAP projections colored by timepoint, showing transcriptional divergence of TCE-induced cells from steady-state populations. Cell numbers per timepoint are indicated. **B**. Absolute CD8⁺ T cell counts per timepoint. **C**. Dot plot of canonical marker genes across the six identified clusters. Dot size indicates the percentage of cells expressing each gene; color intensity indicates scaled mean expression. **D**. UMAP projections colored by cluster identity, resolving six transcriptional states: naïve, memory, progenitor-like, early activated, cycling, and cytotoxic. Cell numbers per cluster are indicated. **E**. Stacked bar plots showing the proportional composition of CD8⁺ clusters at each timepoint. The progenitor-like state dominates at day 3 (54%) before contracting at day 7 (10%), while cytotoxic cells expand from baseline to nearly 30% by day 7. **F**-**G**. Feature plots showing the expression of *Sell* (enriched in naïve and day 3 progenitor-like populations) and *Gzmb* (progressively enriched in cytotoxic cells at day 7) projected onto the UMAP. **H-I: FACS recapitulates TCE-determined evolution of CD8^+^ CTL landscape at the protein level. H**. Experimental design of flow cytometric validation. **I**. Representative plots and quantification of CD62L⁻ granzyme B⁺ CTLs among CD8⁺ T cells at days 0, 3 and 7 post-TCE. Each dot represents one mouse. Data are pooled from 3 independent experiments. P values by one-way ANOVA with Tukey’s adjustment for multiple comparison.

Unsupervised clustering resolved six CD8⁺ T cell states across timepoints (**Figure 4C-E, Extended Data Figure 5A-B**): naïve (*Lef1*, *Sell*, *Ccr7*), memory (*Cd44*, *Il7r*, *Bcl2*), progenitor-like (*Slamf6*, *Tox*, *Tcf7*, *Id3*), early activated (*Nr4a1*, *Bcat1*, *Tnfrsf9*), cycling (*Mki67*, *Ccnb2*), and cytotoxic effector (*Klrg1*, *Gzmb*, *Prf1*). Before treatment, the naïve cluster predominated alongside memory and cytotoxic populations. Notably, the memory cluster co-expressed *Cd44*, *Il7r*, and *Bcl2* together with *Sell*, a profile consistent with central memory T cells (T_CM_) rather than effector memory cells. By day 3, these pre-existing subsets were proportionally diluted by a massive progenitor-like population that robustly expressed *Sell* as well as features of renewal and exhaustion, dominating the CD8⁺ compartment (54% of cells). TCE also induced a prominent early activation program consistent with recent TCR signaling. A cycling cluster (*Mki67, Ccnb2*), detected exclusively at day 3, retained transcriptional features of the progenitor-like and memory states but lacked cytotoxic gene expression, consistent with cell cycle entry from non-effector parent populations. By day 7, activation and proliferation had declined, progenitor-like cells had contracted to 10%, and the cytotoxic subset had doubled to nearly 30% of CD8⁺ T cells. A cytotoxic cluster re-emerged at day 7 that was temporally and transcriptionally distinct from the minor pre-existing cytotoxic population. Because these cytotoxic T lymphocytes (CTL) arose after the progenitor-like transition, we term them CTL_TCE_. Feature plot analysis was consistent with the progressive dynamics of this reprogramming: *Sell* expression was high at day 0 and within the day 3 progenitor-like compartment, before declining by day 7, suggesting transcriptional continuity between pre-existing *Sell*⁺ populations and the progenitor-like state, while *Gzmb* was progressively enriched in CTL_TCE_ at day 7 (**Figure 4F, G**). Protein-level profiling in the spleen of TCE-treated mice corroborated these dynamics, revealing that at day 7, the proportion of CTL (CD62L^-^ granzyme B^+^) had doubled compared to baseline (**Figure 4H, I**).

Together, these data indicate that TCE drives a coordinated, population-level transcriptional reprogramming of the CD8^+^ T cell pool: from pre-existing subsets, TCE induces in 3 days a massive progenitor-like state without lytic potential, followed by an accumulation of CTL_TCE_ defining the biological basis of the recall competence observed at day 7.

### Central memory T cells are the principal source of TCE-induced cytotoxic effectors

We then aimed to characterize which pre-existing CD8^+^ T cell subset displayed the potential to seed CTL_TCE_ via this transcriptional reprogramming. Our single-cell data revealed a transcriptional continuity between the pre-existing memory cluster and the progenitor-like state, both expressing *Sell* (CD62L), before the emergence of CTL_TCE_ at day 7. Trajectory analysis supported a continuous differentiation path from pre-existing populations through the progenitor-like intermediate to cytotoxic effectors (**Figure 5A**), but could not distinguish whether naïve or memory cells initiate this trajectory.

**Figure 5.**
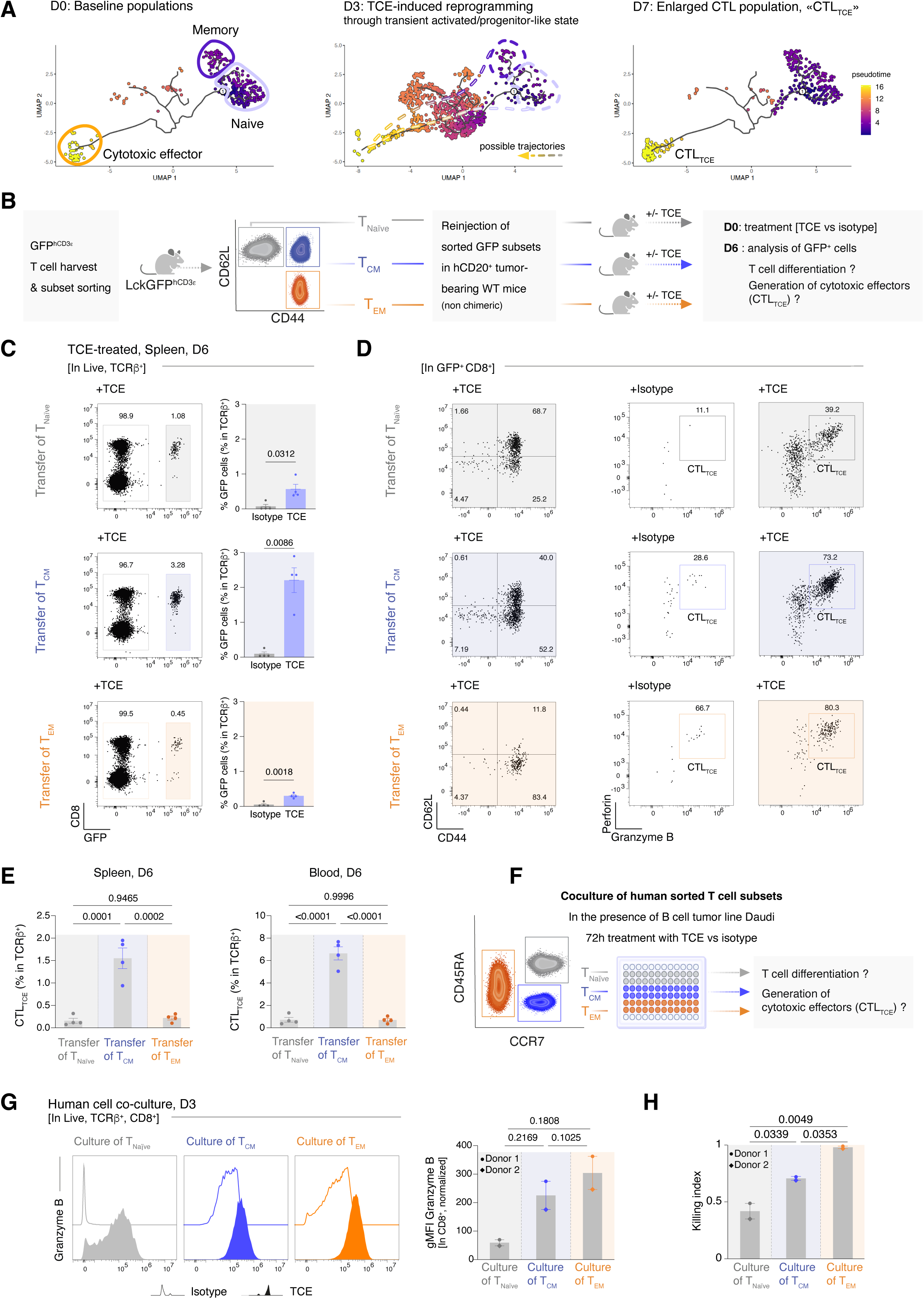
Central memory T cells are the principal source of TCE-induced cytotoxic effectors (CTL_TCE_). **A**. Pseudotime trajectory analysis (Monocle 3) overlaid on the UMAP from Figure 4, suggesting a continuous differentiation path from pre-existing naïve and/or memory populations, through the progenitor-like state, to cytotoxic effectors. **B-E: Murine adoptive transfer reveals that T_CM_ display the highest potential for expansion and differentiation into CTL_TCE_. B**. Experimental design: T_Naïve_ (CD44⁻ CD62L⁺), T_CM_ (CD44⁺ CD62L⁺) and T_EM_ (CD44⁺ CD62L⁻) CD8⁺ T cells are sorted from LckGFP^hCD3ε^ chimeras and transferred into hCD20⁺ tumor-bearing C57BL/6J recipients (non-chimeric) one day before TCE or isotype control administration. GFP⁺ transferred cells are analyzed at day 6 post-treatment. Each dot represents one mouse; data pooled from 2 independent experiments. P values calculated by one-way ANOVA after adjustment using Tukey’s multiple comparisons test. **C-D**: **Characterization of recovered GFP^+^ T cells at D6 from TCE or isotype control**. **From top to bottom: evolution of transferred T_Naïve_, T_CM,_ and T_EM_ subsets upon TCE administration. C.** Representative flow cytometric gating of recovered GFP^+^ T cell subsets and quantification among total T cells, for each transferred subset. **D.** Representative gating of phenotype by CD44 and CD62L expression among recovered GFP^+^ CD8^+^ T cells, and expression of cytotoxic molecules allowing the quantification of CTL_TCE_ (GFP⁺ CD8⁺ perforin^+^ granzyme B⁺). FACSplots from isotype-treated mice are uncolored; TCE-treated FACSplots are colored according to the sorted subset transfer (gray: T_Naïve_; blue: T_CM_; orange: T_EM_). **E**. Quantification of CTL_TCE_ among total T cells depending on subset transfer, in the spleen (left) and the blood (right). **F-H: The superiority of memory subsets over naïve cells for CTL_TCE_ generation is conserved in human co-culture experiments. F**. Experimental design for human validation: T_Naïve_ (CCR7⁺ CD45RA⁺), T_CM_ (CCR7⁺ CD45RA⁻) and T_EM_ (CCR7⁻) CD8⁺ T cells sorted from healthy donor PBMCs are co-cultured with hCD20⁺ B cell tumor in the presence of TCE or isotype control. Analysis is performed after 3 days of co-culture. Two different donors from 2 independent experiments are pooled (Donor 1, circles; Donor 2, diamond). P values by one-way ANOVA with adjustment by Tukey’s multiple comparison test. **G**. Left: Representative histograms of granzyme B expression in CD8⁺ T cells upon treatment with isotype (empty plot) or TCE (full plot), according to the type of subset in culture. Right: Quantification of the gMFI of granzyme B expression in CD8⁺ T cells upon TCE exposure, normalized to the mean gMFI in the isotype control group. **H**. Killing index per subset upon TCE exposure, normalized to the mean killing index in the isotype control group.

To directly test which pre-existing subset seeds CTL_TCE_, we sorted CD8⁺ T cells from LckGFP^hCD3ε^ chimeras into T_Naïve_ (CD62L⁺ CD44⁻), T_CM_ (CD62L⁺ CD44⁺), and T_EM_ (CD62L⁻ CD44⁺) populations and transferred each into tumor-bearing non-chimeric recipients before TCE administration (**Figure 5B**).

The three subsets displayed markedly different responses to TCE (**Figure 5C–D**). T_Naïve_ showed limited expansion and acquired CD44 expression but generated few cytotoxic effectors. T_EM_ retained an effector phenotype and expressed perforin and granzyme B, but barely expanded. T_CM_ combined both capabilities: they showed the highest proliferative expansion and robust acquisition of cytotoxic molecules, with nearly 75% of T_CM_-derived cells co-expressing perforin and granzyme B. As a result, T_CM_ generated substantially more CTL_TCE_ than either T_Naïve_ or T_EM_ across blood and spleen (**Figure 5E**).

To determine whether this hierarchy was conserved in human T cells, we sorted analogous subsets from healthy donor PBMCs (T_Naïve_: CD45RA⁺ CCR7⁺; T_CM_: CD45RA⁻ CCR7⁺; T_EM_: CD45RA⁻ CCR7⁻) and co-cultured them with hCD20⁺ tumor cells in the presence of TCE (**Figure 5F**). Memory-derived cells, both T_CM_ and T_EM_, expressed substantially higher levels of granzyme B than naïve-derived cells and killed tumor cells more efficiently (**Figure 5G–H, Extended Data Figure 6**). The distinction between T_CM_ and T_EM_ was less apparent in this *in vitro* setting than in our *in vivo* transfers, likely because the 72-hour co-culture does not recapitulate the proliferative advantage that distinguishes T_CM_ *in vivo*^34^.

Thus, both murine and human pre-existing subsets display a conserved hierarchy for CTL_TCE_ generation, with memory cells, and T_CM_ in particular, serving as the principal reservoir for expansion and cytotoxic differentiation under TCE.

### TCE induces cDC1-dependent cross-priming and protective tumor-specific memory

Having determined that TCE triggered rapid perforin-dependent cytotoxicity while reprogramming memory T cells to expand the pool of CTLs, we aimed to characterize whether TCE could induce prolonged tumor control after treatment discontinuation. To this end, we subjected tumor-bearing mice to a single injection of TCE or isotype control and followed their survival. While control mice all succumbed rapidly, a majority of TCE-treated mice displayed prolonged, months-long survival (**Figure 6A**), with a survival plateau consistent with the development of an adaptive immune response.

**Figure 6.**
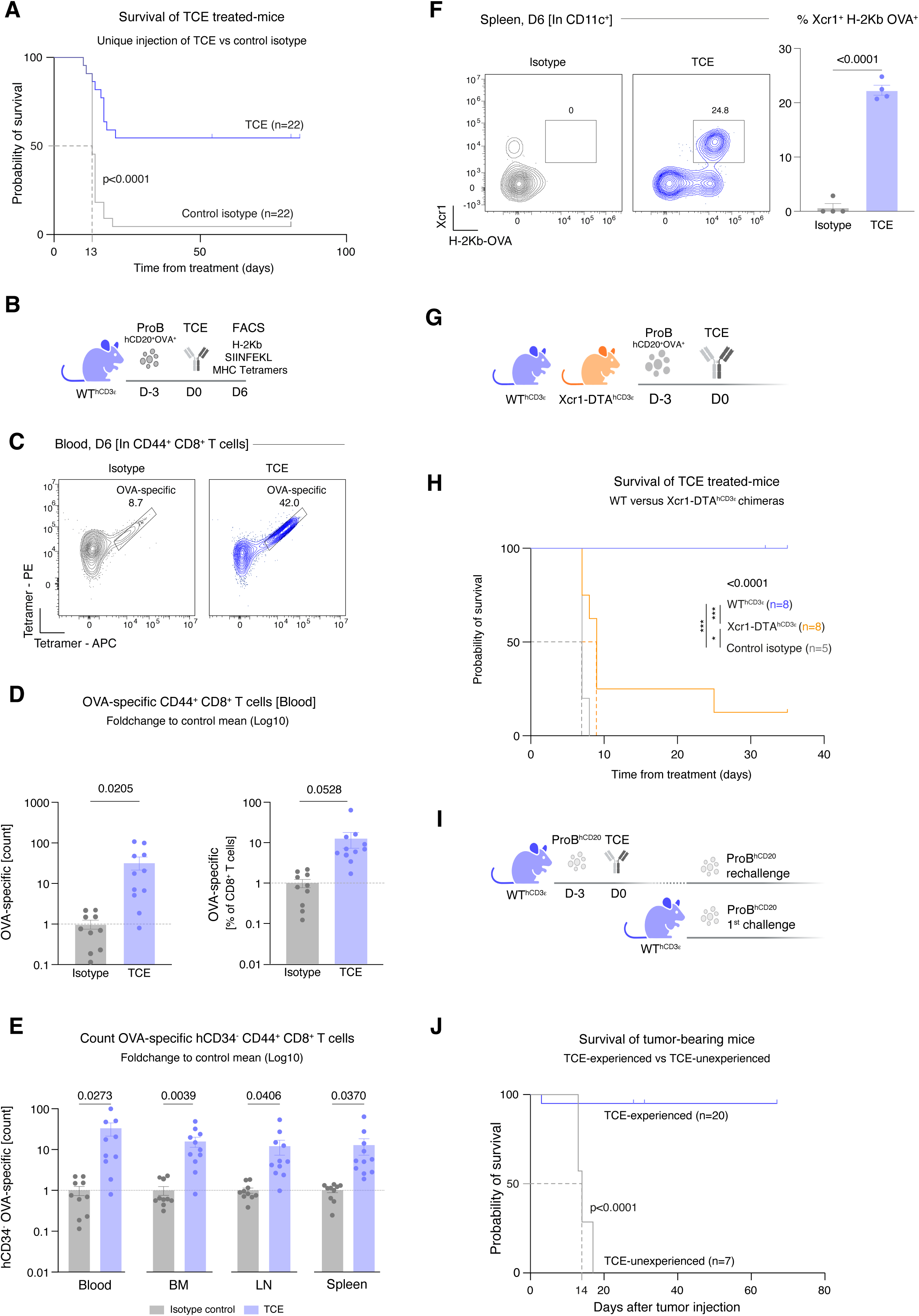
TCE induces cDC1-dependent cross-priming and protective immunological memory. **A**. Kaplan-Meier survival curves of WT^hCD3ε^ mice bearing hCD20⁺ Pro-B lymphoma treated with a single injection of TCE (blue, n = 22) or isotype control (grey, n = 22). Dashed gray line indicates the median survival for control at day 13. P < 0.0001, log-rank test. **B-E: TCE induces de novo priming of antigen-specific CD8^+^ T cells. B**. Experimental design: WT^hCD3ε^ mice bearing hCD20⁺ Pro-B-OVA tumor are treated with TCE or isotype at day 0, and circulating SIINFEKL–H-2Kᵇ tetramer-positive CD8⁺ T cells are analyzed at day 6. **C**-**D**. Representative tetramer staining and quantification of OVA-specific CD44^+^ CD8⁺ T cells (fold-change in proportion and absolute counts, log10 scale). **E**. Foldchange in absolute counts of hCD3ε⁻ tetramer-positive CD44^+^ CD8⁺ T cells across blood, bone marrow, spleen and lymph nodes (log10 scale). Each dot represents one mouse (n=10 isotypes, 11 TCE); data are pooled from three independent experiments. **F-H: cDC1s prime antigen-specific T cells for prolonged antitumor effects of TCE**. **F**. Detection of cross-presented tumor-derived antigen: H-2Kᵇ-OVA complexes on XCR1⁺ cDC1s in TCE-treated versus isotype-treated mice, assessed by flow cytometry (25-D1.16 antibody). **G**. Experimental design: WT^hCD3ε^ and XCR1-DTA^hCD3ε^ chimeras (lacking cDC1s) are established with hCD20⁺ tumor and treated with TCE. **H**. Kaplan-Meier survival curves: WThCD3ε (blue, n = 8) versus XCR1-DTA^hCD3ε^ (orange, n = 8) versus isotype control (gray, n=5). Log-rank test with Holm-Šídák’s multiple comparisons test. **I-J: TCE promotes protective adaptive antitumor immunity**. **I.** Rechallenge experimental design: TCE-unexperienced chimeras versus mice cured by a single TCE injection (TCE-experienced) are (re)injected with hCD20⁺ Pro-B tumor without any further treatment. **J**. Kaplan-Meier survival curves after rechallenge: TCE-experienced (blue, n = 20) versus TCE-naïve (grey, n = 7). P < 0.0001, log-rank test.

To determine whether TCE could generate tumor-specific responses, we introduced the model antigen OVA into our hCD20⁺ Pro-B tumor cell line (**Figure 6B**), allowing us to monitor the generation of SIINFEKL-H-2Kᵇ tetramer-positive tumor-specific CD8⁺ T cells. At day 6 from treatment, circulating tetramer-positive CD8⁺ T cells were specifically enriched in TCE-treated mice, with an approximately 30-fold increase in absolute counts (**Figure 6C, D**).

To determine whether this expansion only relied on direct TCE-mediated activation, we took advantage of the incomplete chimerism of our model: hCD3ε-negative T cells, which cannot be engaged by TCE, thus serve as a built-in internal control for *de novo* priming independent of TCE engagement. Tetramer-positive cells were robustly enriched among hCD3ε-negative T cells across all organs examined (**Figure 6E**). These results demonstrate that TCE-induced killing triggers *bona fide de novo* priming of tumor-specific T cells independently of TCE engagement.

We hypothesized that cross-presenting type 1 conventional dendritic cells (cDC1s) were responsible for this *de novo* priming. Consistent with this, XCR1⁺ cDC1s upregulated expression of H-2Kᵇ-OVA complexes in TCE-treated mice (**Figure 6F**), directly demonstrating cross-presentation of tumor-derived antigens following TCE-induced killing. To test the functional requirement for cDC1s, we generated XCR1-DTA^hCD3ε^ chimeras lacking cDC1s alongside WT^hCD3ε^ controls and treated tumor-bearing animals with TCE (**Figure 6G**). In cDC1-deficient chimeras, tumor-specific T cells were undetectable (**Extended Data Figure 7**). Crucially, while WT chimeras achieved durable tumor control, nearly all cDC1-deficient mice succumbed despite identical TCE treatment (**Figure 6H**), establishing that cDC1-mediated cross-priming is essential for the long-term therapeutic benefit of TCE.

Finally, to investigate whether TCE could confer lasting protective immunity against endogenous tumor antigens, we rechallenged mice that had been cured from (OVA-negative) hCD20⁺ Pro-B lymphoma by a single injection of TCE with the same tumor line, without any further treatment, alongside naïve chimeras as controls (**Figure 6I**). While TCE-naïve mice all succumbed within two weeks of tumor injection, the vast majority of TCE-experienced mice remained tumor-free after rechallenge (**Figure 6J**).

Together, these results demonstrate that despite bypassing TCR-pMHC recognition, TCE-mediated killing generates the antigenic substrate for cDC1-dependent cross-priming of tumor-specific CD8⁺ T cells, establishing durable immunological memory that enables tumor control independently of continued treatment. T cell engagers thus function not merely as a transient pharmacological bridge between T cells and the tumor, but also as initiators of a self-sustaining adaptive immune response.

## DISCUSSION

In this study, we developed an immunocompetent chimeric mouse model to dissect the mechanism of action of human T cell engagers *in vivo*. By combining intravital imaging, single-cell transcriptomics, adoptive transfers, and genetic loss-of-function approaches, we reveal that TCE activity unfolds through temporally distinct phases: rapid perforin-dependent tumor killing, a refractory period of active T cell reprogramming, and the establishment of cDC1-dependent tumor-specific adaptive immunity. This sequence demonstrates that TCEs, designed to bypass TCR-pMHC recognition, ultimately restore it, coupling transient pharmacological cytotoxicity to durable immunological memory.

The WT^hCD3ε^ model addresses a longstanding gap in the preclinical study of TCEs. Unlike humanized NSG mice reconstituted with human PBMCs, which develop graft-versus-host disease and lack a complete immune microenvironment^27^, WT^hCD3ε^ chimeras maintain physiological T cell development and differentiation, and normal immune regulation, while enabling the direct evaluation of clinical-grade human TCEs. Previous approaches to humanize the CD3ε locus have generated valuable tools^28,29^ but have not been combined with the genetic versatility of bone marrow chimeras, which allowed us to interrogate specific effector pathways through *Prf1*KO, *Ifng*KO, and *XCR1*-DTA chimeras within a unified experimental framework.

Our intravital imaging data reveal an unexpected temporal dissociation between molecular engagement and killing competence. While TCE establishes stable T cell-tumor conjugates within minutes, around 3% of these contacts were lytic during the first hours, rising to almost 30% by 24 hours. This delayed acquisition of cytotoxic function likely reflects the time required for granule polarization and effector molecule upregulation^35,36^, and may rely on the activation of pre-existing CTLs that acquire lytic competence within a day. The absolute dependence on perforin, consistent with death receptor-independent killing^20^, and on LFA-1, first identified three decades ago^37^ and subsequently confirmed in structural studies^38,39^, indicates that TCEs kill through integrin-stabilized contacts. Our intravital data extend these findings by demonstrating the kinetics and functional consequences of integrin-dependent contact formation *in vivo*.

A central finding is the biphasic cytotoxic response to TCE: an acute killing wave at day 1, a refractory period at days 2-3 during which abundant T cells fail to kill tumor cells despite re-exposure to TCE, and the emergence of immediate recall cytotoxicity at day 7. Our single-cell data indicate that this temporal pattern reflects the sequential mobilization of functionally distinct CD8⁺ T cell populations rather than a uniform loss and recovery of effector function.

At baseline, a minor pre-existing cluster of cytotoxic CD8⁺ T cells expresses the effector molecules required for immediate killing. These cells bear hallmarks of terminally differentiated short-lived effectors (*Zeb2, Klrg1, Cx3cr1*)^40,41^, and their transcriptional profile overlaps with that of FACS-defined effector memory T cells, consistent with the limited expansion potential observed upon T_EM_ transfer in our adoptive transfer experiments. By day 3, this small effector population is numerically overwhelmed by a large TCE-induced compartment that lacks immediate cytolytic capacity, which coincides with the collapse of killing activity and may account for the population-level unresponsiveness to TCE re-injection.

The dominant TCE-induced population at day 3 is transcriptionally distinct from all pre-existing subsets. It co-expresses *Tcf7, Slamf6,* and *Id3*, a molecular signature characteristic of precursors of exhausted T cells (T_pex_)^42–45^. Strikingly, these progenitor-like cells retain expression of *Sell* (CD62L), a feature they share with the pre-existing memory cluster, which itself co-expresses *Sell*, *Cd44*, *Il7r*, and *Bcl2*, a profile corresponding to central memory T cells (T_CM_). This transcriptional continuity, from a *Sell*⁺ memory state through a *Sell*⁺ progenitor-like intermediate, pointed to T_CM_ as the primary source of the reprogramming wave. Adoptive transfer of sorted subsets directly confirmed this trajectory: T_CM_ displayed the highest expansion and the most robust acquisition of cytotoxic molecules, generating substantially more CTL_TCE_ than either naïve or effector memory T cells. In human co-culture experiments, the superiority of memory-derived cells over naïve cells for granzyme B expression and tumor killing was conserved across donors and TCE formats. The distinction between T_CM_ and T_EM_ was less apparent in this *in vitro* setting, likely because the 72-hour co-culture does not recapitulate the proliferative advantage that distinguishes T_CM_ from T_EM_ *in vivo*^34^. Together, these findings identify the pre-existing central memory compartment as the principal reservoir for CTL_TCE_ generation, resonating with CAR-T cell data linking memory-phenotype cells to superior clinical outcomes^46–48^, and with recent clinical observations associating pre-existing T cell composition with TCE response^14,24,26^.

By day 7, the progenitor-like compartment had contracted while CTL_TCE_ expanded to approximately 30% of CD8⁺ T cells, coinciding with restored cytotoxicity. Several non-exclusive mechanisms may contribute to the refractory period itself, including target antigen downregulation^49^, TCE consumption^50^, or functional exhaustion amenable to treatment-free intervals^51^. However, our data suggest that such intervals serve a purpose beyond passive recovery: they provide the temporal window required for T_CM_-derived T cells to transit through the progenitor-like intermediate and differentiate into CTL_TCE_ endowed with durable cytotoxic function.

Why does a T_pex_-like program, typically associated with weeks of chronic antigen exposure^42–45^, arise within days of a single TCE injection? We propose that this reflects the signaling imbalance inherent to TCE-mediated activation. TCE delivers a strong, sustained signal through CD3ε crosslinking that bypasses TCR-pMHC selectivity^19^. In a conventional immune synapse, this proximal signal is matched by costimulation from professional antigen-presenting cells. In the TCE context, even when tumor cells express costimulatory molecules, as B-lineage malignancies often do, the balance between CD3-driven NFAT activation and costimulation-driven AP-1 engagement is tilted toward unopposed NFAT signaling^52^, a configuration that drives *Tox* expression and the progenitor/exhaustion transcriptional program^53–55^. Consistent with this signaling model, a recent clinical study of glofitamab in large B cell lymphoma demonstrated that providing exogenous 4-1BB costimulation alongside TCE shifted intratumoral T cells toward a naïve-like state and enhanced antitumor efficacy^14^, suggesting that insufficient costimulation may underlie the progenitor-like transition. Whether *Tox* co-expression within this progenitor-like compartment marks a bifurcation toward terminal exhaustion rather than productive differentiation will require epigenetic profiling to resolve^54,55^, a distinction with direct implications for TCE/anti-PD-1 combinations^56^.

Perhaps the most significant finding is the demonstration that TCE-mediated killing provides the antigenic substrate for a cDC1-dependent cross-priming cascade that generates tumor-specific adaptive immunity. The expansion of antigen-specific T cells among hCD3ε-negative cells, which cannot be engaged by TCE, directly demonstrates *de novo* priming through conventional antigen presentation, supported by the loading of tumor-derived peptide–MHC complexes on XCR1⁺ cDC1s and the loss of durable tumor control in cDC1-deficient mice. TCEs have been designed to operate outside the endogenous cancer-immunity cycle^57^, killing tumors without requiring its upstream steps. Our data show that this killing feeds back into this cycle: tumor lysis provides the antigenic substrate, cDC1 cross-presentation converts it into T cell priming, and protective memory follows. What distinguishes the TCE context from other cytotoxic modalities is the concurrent polyclonal T cell activation it triggers: perforin generates the antigenic substrate while IFN-γ may license dendritic cells for cross-presentation, as described in checkpoint blockade settings^58^. Whether TCE-mediated killing formally constitutes immunogenic cell death, involving calreticulin exposure, ATP release, HMGB1 secretion, or activation of the cGAS-STING axis by tumor-derived DNA^59^, remains to be determined. Our data establish the functional consequence of immunogenic killing without resolving the molecular intermediates, an important area for future investigation that may reveal additional targets for combination strategies.

The survival plateau and the protective memory demonstrated by rechallenge experiments together provide a biological framework for limited-duration TCE therapy^50^. These findings are consistent with recent reports of rechallenge protection in immunocompetent glioma models, where bispecific antibody-treated mice rejected antigen-loss tumor variants, suggestive of epitope spreading^15^. Our data provide a mechanistic basis for these observations: the cDC1-dependent cross-priming cascade we identify converts TCE-mediated killing into antigen-specific memory that persists independently of continued treatment. Once this cascade has been initiated, continued administration may be unnecessary, consistent with durable remissions after discontinuation of mosunetuzumab and glofitamab^10,60^. From a translational perspective, our findings argue for preserving the T_CM_ compartment before treatment, as it serves as both the principal effector reservoir and a potential predictive biomarker of response. They further support combinations that preserve innate responses^61^ and amplify the cross-priming axis. These could include Flt3L or STING agonists for cDC1 function^62–65^, checkpoint blockade to expand progenitor-like populations^42,44^, and vaccines to reinforce tumor-specific priming^66^. In addition, they argue for the monitoring of tumor-specific T cell responses as biomarkers of durable remission.

Several limitations warrant consideration. Our model employs murine TCRs, an aggressive and transplanted Pro-B tumor, and a strong model antigen (OVA), which may overestimate cross-priming efficiency. The hCD3ε transgene is overexpressed, potentially affecting TCE binding kinetics and signaling intensity^19,67^. Chimera generation requires lethal irradiation of recipients and can result in incomplete chimerism, which, while providing an informative internal control, means a fraction of T cells are not engaged by TCE; however, clinical pharmacokinetics create analogous drug-free windows^50^. The roles of CD4⁺ T cells and IFN-γ in sustaining cross-priming were not directly tested. The applicability to solid tumors, where the microenvironment differs fundamentally^68,69^, remains to be determined. Furthermore, our *in vitro* human experiments used the Daudi Burkitt lymphoma line, which may not represent the diversity of B cell malignancies encountered clinically.

In conclusion, we reveal that T cell engagers orchestrate a coordinated immune program extending far beyond polyclonal T cell redirection. Perforin-dependent killing by pre-existing effectors, transcriptional reprogramming of central memory T cells into CTL_TCE_ through a progenitor-like intermediate, and cDC1-mediated cross-priming collectively couple the destruction of tumor cells to the construction of lasting adaptive immunity against them.

## Extended Data Figure legends

**Extended Data Figure 1 | Normal T cell development and homeostasis in WT^hCD3ε^ mice.**

**A.** Representative flow cytometry plots and quantification of thymic T cell subsets (CD4 SP, CD8 SP, DP, DN) gated on live, lineage-negative (CD11b⁻ TCRγδ⁻ NK1.1⁻ CD11c⁻ B220⁻) cells in WT and WT^hCD3ε^ mice. **B.** Representative flow cytometry plots of CD4⁺ and CD8⁺ T cell compartments in lymph nodes (LN, top) and spleen (bottom) from WT and WT^hCD3ε^ mice, showing CD4/CD8 ratios (left) and CD44/CD62L-defined T_Naïve_, T_CM_, and T_EM_ subsets within CD8⁺ (center) and CD4⁺ (right) populations. **C.** Quantification of T cell subset proportions (normalized to WT mean) in LN (top) and spleen (bottom) for CD4⁺ and CD8⁺ compartments. **D.** Representative flow cytometry plots of FoxP3⁺CD25⁺ regulatory T cells among CD4⁺ T cells in LN (top) and spleen (bottom). **E.** Quantification of Treg frequency (% of CD4⁺, normalized to WT mean) in blood, LN, and spleen. **F.** hCD34 reporter expression (as a surrogate of hCD3ε expression, % among TCRβ⁺ cells) in peripheral blood across three representative chimera screening batches. Each dot represents one mouse from a single experiment. P values by two-tailed unpaired t-test between WT and WT^hCD3ε^ for each subset independently.

**Extended Data Figure 2 | TCE induces T cell activation *in vitro* against hCD20⁺ target cells.**

**A.** Percentage of CD44⁺ cells among CD8⁺ hCD34⁺ T cells after co-culture with hCD20⁺ Pro-B cells (20 PBMC: 1 tumor cell) in the presence of isotype control, CD20-TCE, mosunetuzumab (Mosun), or glofitamab (Glofi). **B.** Left: percentage of CD25⁺CD69⁺ cells among CD4⁺ hCD34⁺ T cells co-cultured with hCD20⁺ Pro-B cells. Right: percentage of CD44⁺ cells in the same CD4⁺ compartment upon treatment with isotype, TCE, mosunetuzumab, or glofitamab. **C.** Percentage of PD-1⁺CD69⁺ (left) and CD44⁺ (right) cells among CD8⁺ hCD34⁺ T cells after co-culture with hCD20⁺ Eμ-Myc tumor cells. **D.** Same as (**C**) for CD4⁺ hCD34⁺ T cells. Each dot represents one well from a single experiment, except for CD25⁺CD69⁺ quantification in (**B**, left), where dots represent means of technical replicates pooled from three independent experiments. P values by two-tailed unpaired t-test with Welch’s correction (two-group comparisons) or by paired t-test (**B**, left, pooled data).

**Extended Data Figure 3 | TCE triggers tumor cell apoptosis and perforin–granzyme-dependent killing *in vivo*.**

**A.** Schematic of the bone marrow (BM) chimera strategy for intravital imaging: BM from Lck-Cre × Rosa-mTmG donors is harvested, transduced with hCD3ε, and transplanted into lethally irradiated WT recipients to generate LckGFP^hCD3ε^ chimeras. **B.** Representative flow cytometry plots and quantification of apoptotic tumor cells in the spleen at day 1 after isotype control or CD20-TCE treatment. Apoptosis reporter corresponds to a derived parameter using the ratio of cyan fluorescent protein (CFP) to FRET fluorescence in tumor cells expressing the caspase-3-sensitive CFP-DEVD-YFP reporter. **C.** Percentage of Granzyme B⁺ Perforin⁺ cells among CD4⁺ hCD34⁺ T cells in BM and spleen. **D.** Percentage of CD107a⁺ (degranulation marker) cells among CD8⁺ hCD34⁺ T cells after 6 h of co-culture with or without hCD20⁺ Pro-B target cells and TCE. **E.** Representative flow cytometry plots and quantification of Perforin⁺ Granzyme B⁺ CD8⁺ T cells in the spleen at day 3 after treatment with isotype, TCE, or TCE combined with anti-LFA-1 blocking antibody. In (**B**), (**C**), and (**E**), each dot represents one mouse from a single experiment. In (**D**), each dot represents one well from a single *in vitro* experiment. P values by two-tailed unpaired t-test with Welch’s correction (two-group comparisons) or by one-way ANOVA with Tukey’s adjustment for multiple comparisons (more than two groups).

**Extended Data Figure 4 | TCE treatment remodels the chemokine landscape in spleen and bone marrow.**

**A.** Concentrations (pg/ml) of the indicated chemokines (CXCL10, CCL2, CCL7, CCL3, CCL4, CXCL2, CCL5, CCL11, CXCL1) measured by multiplex ELISA in spleen lysates of tumor-bearing mice treated with isotype control, or TCE at day 1 (D1) and day 3 (D3). **B.** Same as (**A**) for BM lysates. P values by one-way ANOVA with Tukey’s adjustment for multiple comparisons.

**Extended Data Figure 5 | Transcriptomic landscape of CD8⁺ T cell states during TCE treatment.**

**A.** Heatmap of differentially expressed genes across CD8⁺ T cell clusters identified by single-cell RNA sequencing: naïve, memory, progenitor-like, early activated, cycling, and cytotoxic effector. Inset: stacked bar plots showing the absolute count and proportional composition of CD8⁺ T cell clusters at D0, D3, and D7.

**Extended Data Figure 6 | Sorting strategy and human validation of subset hierarchy.**

**A.** Representative flow cytometry plots showing the sorting strategy and post-sort purity of T_Naïve_ (CD62L⁺ CD44^-^), T_CM_ (CD62L⁺ CD44^+^), and T_EM_ (CD62L^-^ CD44^+^) CD8⁺ T cells from hCD34-enriched spleen and LN cells of LckGFP^hCD^^3^^ε^ chimeras. **Human co-culture**. **B.** Left: Representative histograms of Granzyme B expression in CD8^+^ T cells after 3 days of co-culture between CD8⁺ T cells sorted from healthy donor PBMCs (T_Naïve_ (CCR7⁺ CD45RA⁺), T_CM_ (CCR7⁺ CD45RA⁻) and T_EM_ (CCR7⁻)) and hCD20⁺ B Daudi tumor cells, treated with isotype (open histogram), TCE (light filled histogram), or glofitamab (dark filled histogram). Right: quantification of Granzyme B geometric mean fluorescence intensity (gMFI) in CD8⁺ T cells, for each subset co-culture condition, normalized to the isotype control mean. **C.** For each donor, the remaining tumor load in each well was calculated and normalized to the mean tumor load in the [T_Naïve_ transfer + isotype control] condition. The killing index was defined for each culture condition as 1-[normalized tumor load]. Two donors from two independent experiments are pooled (Donor 1, circles; Donor 2, diamonds). P values by one-way ANOVA with Tukey’s adjustment for multiple comparisons.

**Extended Data Figure 7 | cDC1 cells are required for TCE-induced tumor-specific CD8⁺ T cell priming.**

Percentage of OVA-specific CD44⁺ cells among CD8⁺ T cells in WT^hCD3ε^ and Xcr1-DTA^hCD3ε^ (cDC1-depleted) mice 6 days after TCE treatment. Each dot represents one mouse, from one experiment. P v

**Extended Data Video 1 | Before CD20-TCE injection, T cells are motile and colocalize with tumor cells without engaging in stable contacts**

Representative intravital video of one BM position, pre-TCE injection: T cells (green) appeared motile and engaged in transient contacts with tumor cells (gray/white), with a median duration of 7 minutes.

**Extended Data Video 2 | T cells establish stable hourlong interactions with tumor cells within minutes of CD20-TCE injection on D0**

Representative intravital video of one BM position, post-TCE injection: T cells (green) establish stable contacts with tumor cells (gray/white), many of which were still ongoing hours after TCE injection.

**Extended Data Video 3 | Killing is ongoing at D1 after CD20-TCE injection**

Representative intravital video of one BM position, 1 day after TCE injection. Lysis of tumor cells (live (gray/white) to apoptotic (blue)) by T cells (green) was markedly enhanced at 24 hours post-injection, at which point almost 30% of T cell contacts were lytic.

**Extended Data Video 4 | T cells are proliferating at the tumor site on D2 after CD20-TCE injection**

Representative intravital video of one BM position, on day 2 after TCE injection. White squares outline examples of mitosis of T cells (green), among the tumor mass (live (gray/white); apoptotic (blue)).

**Extended Data Video 5 | T cells display immediate recall cytotoxicity when TCE is reinjected at D7**

Representative intravital video of one BM position, on day 7 after TCE injection. Administration of a second dose of TCE triggered immediate tumor killing ((live (gray/white); apoptotic (blue)), suggesting that T cells (green) had acquired enhanced cytotoxic competence during the refractory period.

## MATERIALS AND METHODS

### CELL LINES

#### Murine tumors

Murine B-cell lymphoma lines (Eμ-Myc-derived or Pro-B) were transduced to express the human CD20 molecule (NM_021950.4, human CD20), as well as a FRET-based reporter of caspase-3 activity (pMSCV-CFP-DEVD-YFP)^31^ to evaluate induction of apoptosis in tumor cells. For the exploration of antigen-specific immune responses, hCD20^+^DEVD^+^ Pro-B tumor cells were transduced with ovalbumin (pMSCV-cOVA), and SIINFEKL presentation was confirmed by H2kb-OVA_257-264_ staining (Addgene #25097)). **Human tumors:** The human Burkitt-like B cell lymphoma line Daudi was obtained from Olivier Schwartz’s lab in Institut Pasteur. Cells were retrovirally transduced to express the FRET-based reporter of caspase-3 activity. **Human blood samples and PBMC isolation:** Peripheral blood was obtained from healthy adult donors through the Etablissement Français du Sang (EFS, Île-de-France, n°CPSL 2025-2028-001). All donors provided written informed consent in accordance with the Declaration of Helsinki. Sample collection and processing were approved by the Comité de Protection des Personnes (CPP Île-de-France, approval n°HS_2022_24408) and the Institut Pasteur Ethics Committee. Human CD3 cells were secondarily enriched through magnetic labeling. **Cell culture:** All cell lines were cultured in RPMI supplemented with 10% heat-inactivated fetal bovine serum, penicillin (50 U ml^−1^), streptomycin (50 µg ml^−1^), 1 mM sodium pyruvate, 10 mM Hepes, and 50 µM 2-mercaptoethanol and maintained at 37°C and 5% CO2. Cell lines were routinely tested for the absence of mycoplasma contamination (Venor-GeM Advance Mycoplasma Detection Kit, Minerva Biolabs).

### PLASMIDS

#### Packaging vectors & transduction protocol

We used commercial pCL-Eco (Novus Biotech cat. NBP2-29540) and pCL-Ampho (Novus Biotech cat. NBP2-29541) that were initially described by Naviaux et al^70^ in 1996. Viral particles were produced in HEK293T cells and supernatants were collected before spin infection of target cells (centrifugation at 1000g, 1h30, 32°C). **pMSCV-humanCD3-T2A-eGFP and pMSCV-humanCD3-T2A-humanCD34**: Human CD3 epsilon coding sequence (NM_000733.3) was associated with a T2A self-cleaving sequence and alternatively to the eGFP coding sequence or to a truncated human CD34 coding sequence, synthesized and subcloned into pMSCV backbone. **pMSCV-humanCD20**: Human membrane spanning 4-domains A1 transcript variant 3 coding sequence (NM_021950.4, human CD20) was synthetized and subcloned into pMSCV backbone. **pMSCV-cOVA**: cytosolic chicken ovalbumin coding sequence (cOVA) was subcloned from pCI-neo-cOVA (Addgene #25097) into pMSCV backbone. **pMSCV-CFP-DEVD-YFP**: Our reporter for caspase 3 activity (so-called DEVD) was previously described in Breart et al^31^ in 2008. The initial coding sequence was subcloned from pcDNA3.1 into pMSCV backbone.

### MICE

#### Mouse strains

Six to eight-week-old male C57BL/6J mice were obtained from Envigo. *Ifng*^−/−^, *Prf1*^−/−^, Lck-cre x ROSA-mTmG, and XCR1-cre x Rosa-DTA mice (all under a C57BL/6 background) were bred in our animal facility under specific-pathogen-free conditions, air renewal, ambient temperature of 22 ± 2 °C and controlled light–dark cycle. All animal experiments were conducted with approval from the Institut Pasteur Safety Committee, in compliance with French and European guidelines (CETEA 230038) and were validated by the French Ministry of Education and Research (APAFIS #45558-2023103115139255 v2). **BM chimera generation:** On the 1^st^ day, bone marrow was harvested from femurs and tibias of donor mice (WT, *Ifng*^−/−^, *Prf1*^−/−^, Lck-cre x ROSA-mTmG, or XCr1-cre x Rosa-DTA). Red blood cells were removed through Ficoll-Paque separation and hematopoietic cells were isolated (Corning). Cells were then washed and resuspended in complete media supplemented with 100ng of murine Stem Cell Factors, 20ng of m-IL-3 and 20ng of m-IL-6 (Peprotech) and incubated overnight at 37°C. On days 2 and 3, cells were retrovirally transduced to express the hCD3ε-T2A-hCD34 construct (2 spin infections, 24 hours apart). On day 3, wild type C57BL/6J recipient mice were irradiated with two 5,5 Gy doses, 4 hours apart. Immediately after the 2^nd^ session of irradiation, mice were reconstituted with a total of 3 to 5 × 10^6^ transduced bone marrow cells via intravenous injection. Eight to twelve weeks after reconstitution, mice were bled to evaluate expression of hCD3ε and the transduction marker hCD34 on T cell and non-T cell lineages. Human CD34 was expressed by a majority of BM-derived immune cells, and hCD3ε expression was restricted to mCD3ε^+^ cells. As the antibodies against hCD3ε used in FACS stainings recognize the same epitope as CD20-TCEs, hCD34 expression in T cells was used as a surrogate for hCD3ε expression. The resulting BM chimeras are subsequently referred to as WT^hCD3ε^, LckGFP^hCD3ε^, IfngKO^hCD3ε^, Prf1KO^hCD3ε^, and Xcr1-DTA^hCD3ε^.

### TREATMENTS & INJECTABLE REAGENTS

#### CD20-T cell engagers

The following human CD20-T cell engagers were used for murine and human experiments: research-grade CD20-TCE^8^ was obtained from Genentech under Material Transfer Agreement, clinical grade Mosunetuzumab and Glofitamab were purchased from Cliniscience. As a control, the human IgG1 PG-LALA isotype was purchased from BioXcell. **Anti-LFA-1 antibody:** For LFA-1 blockade, the anti-CD11a (clone M17/4) was obtained from BioXcell. As a control, a rat IgG2a_k_ (clone 2A3) isotype from BioXcell was used. All antibodies were injected IV (in 200μl PBS), at a dose of 1mg.kg^-1^.

### IN VITRO EXPERIMENTAL SETUP

#### Murine cytotoxicity assays

T cell frequencies in PBMC preparations collected from WT^hCD3ε^ chimeras were assessed by flow cytometry before each experiment and represented 28.4 ± 3.6% of live cells (mean ± SEM, N = 3 independent experiments, total of 24 mice). Based on this proportion (∼25–30%), we plated 20 PBMCs per target cell as a proxy for an E:T ratio of approximately 5:1. Tumor targets were hCD20-expressing murine B cell tumors (hCD20^+^DEVD^+^ Pro-B or hCD20^+^ Eμ-Myc cells), co-cultured for 48 to 72 hours in 96-well round-bottom plates, in the presence of 100ng/ml research-grade CD20-TCE or human IgG1 LALA-PG isotype control. The TCE concentration of 100 ng/mL was selected based on the publication first describing research-grade CD20-TCE^8^. T cell activation was assessed by flow cytometry. **Human cytotoxicity assays:** Human CD3^+^ T cells purified from PBMC (Pan T Cells Isolation Kit human, from Miltenyi) were co-cultured with Daudi tumor cells at an E:T ratio of 5:1, 96-well plates for 72 hours, in presence of 100ng/ml CD20-TCE (research- or clinical-grade) or human IgG1 LALA-PG isotype control. For each donor, the remaining tumor load in each well was calculated and normalized to the mean tumor load in the [T_Naïve_ transfer + isotype control] condition. The killing index was defined for each culture condition as 1-[normalized tumor load].

### IN VIVO EXPERIMENTAL SETUP

#### Tumor implantation

Three days before treatment administration, recipient mice received 2 or 10×10^6^ hCD20^+^DEVD^+^ Pro-B or OVA^+^hCD20^+^DEVD^+^ Pro-B tumor cells as specified in figure legends. All murine tumor cells were injected IV in 200 μl PBS. ***In vivo* treatment with CD20-TCE:** When appropriate, anti-LFA-1 was injected IV one day before TCE administration. At day 0, mice were treated with 1mg.kg^-1^ CD20-TCE or human IgG1 LALA-PG isotype control. For recall experiments, a second dose of TCE was administered at defined timepoints (e.g., day 3, day 7) during imaging sessions.

### READOUTS & TECHNOLOGIES

#### Detection of tumor apoptosis in real-time

As previously described^31^, tumor cells are transduced with the CFP-DEVD-YFP construct, allowing for emission in the YFP wavelength upon CFP excitation via FRET (Fluorescence Resonance Energy Transfer). Upon caspase-3 activation, DEVD is cleaved, and the loss of FRET can be detected via FACS or 2-photon intravital imaging.

### TCE-induced T cell transformation by flow cytometry

Bone marrow (flushed tibia & fibula), spleen and lymph nodes (gentle dissociation and enzyme digestion when DCs were evaluated), and blood (RBC lysis) were processed into single-cell suspensions. **Murine cell surface staining**: Cells were incubated in FACS buffer (2% Fetal Bovine Serum or FBS, 0.2% EDTA in PBS) with anti-CD16/ CD32 antibody (clone93, Biolegend), fixable viability dye eF780 (1:1000, Invitrogen), and the following antibodies for 20 min at 4°C. hCD34-APC (clone 561, BioLegend), hCD3-PE (Clone UCHT1, BioLegend), CD45.2-BUV737(clone 104, BD Biosciences), TCRβ-PE-Cy7(clone H57-597, BioLegend), CD4-BUV395 (clone GK1.5, BD Bioscience), CD8α-BV785 (clone 53-6.7, BioLegend), CD44-BV605 (clone IM7, BioLegend), CD62L-BV421 (clone MEL-14, BioLegend). CD45.1-BV650 (clone A20, BioLegend), CD45.2-BV650 (clone 104, BioLegend), hCD20-BUV661 (clone 2H7, BD Biosciences). CD69-BUV661(clone H1.2F3, BD-Biosciences), CD25-PE (clone PC61, BioLegend), PD-1-PECy7 clone (29F.1A12, BioLegend), LFA-1-APC (clone H155-78, BioLegend). H-2Kb APC or PE (clone alexa6-88.5, BioLegend), H-2Kb-SIINFELK OVA_257-264_ APC or PE (clone 25-D1.16, BioLegend), IA-IE-BUV395 (clone M5/114.15.2, BD Biosciences), CD86-BV650 (clone GL-1, BioLegend), CD11c-APC or PE (clone N418 or HL3, BioLegend), XCr-1-BV785 (clone ZET, BioLegend). CD4-BUV395 (clone GK1.5, BD Biosciences), CD62L-BV421 (clone MEL-14, BioLegend), CD44-BV605 (clone IM7, BioLegend)), CD45.1-BV650 (clone A20, BioLegend), CD8a-BV785 (clone 53-6.7, BioLegend)), Granzyme B-PE (clone QA18A28, BioLegend), TCRb-PECy7 (clone HTK888, BioLegend), Perforin-APC (clone S16009B, BioLegend). hCD34-APC (clone 561, BioLegend), CD44-BV605 (clone IM7, BioLegend) and CD62L-BV421 (clone MEL-14, BioLegend). For evaluation of OVA-specific CD8^+^ T cell responses, H2-Kb-OVA_257-264_ MHC:peptide tetramers were kindly provided by the National Health Institute Tetramer Core Facility. Tetramer staining was conducted first at 4°C for 1 hour in a dedicated buffer (2% Fetal Bovine Serum or FBS, 0.2% EDTA, 0,1% NaN3 in PBS), followed by staining with the antibody cocktail. For the degranulation assay, cells were resuspended in R10 in the presence of monensin and brefeldin A (BD Biosciences) and stained with CD107a-PE (clone 1D4B, BioLegend) for 4 hours at 37°C. **Intracytoplasmic staining:** After extracellular staining, cells were fixed and permeabilized according to the manufacturer’s protocol (Cytofix/cytoperm kit from BD) and stained with the following antibodies: Granzyme B-PE (clone QA18A28), Perforin-APC (clone S16009B). **Intranuclear staining**: After extracellular staining, cells were fixed and permeabilized according to manufacturer’s protocol (Transcription Factor Staining Buffer from Invitrogen cat 00-5523-00) and stained with the following antibodies: Ki67-BV421 (clone 16A8, BioLegend), FoxP3-PE (clone FJK-16s, BioLegend). **Human cell staining**: For standard human cell surface staining, the cells were incubated in FACS buffer (2% FBS, 0.2% EDTA in PBS) with Human TruStain FcX (BioLegend), fixable viability dye eF780 (1:1000, Invitrogen), and the following antibodies for 20 min at 4°C: CD8-BV421 (clone SK1, BioLegend), CD4 SK3-BV785 (clone OKT4, BioLegend), CD19-BV650 (clone SJ25C1, BioLegend)), CD27-BV605 (clone O323, BioLegend), CCR7-PE (clone G043H7, BioLegend), CD45RO-BUV661(clone UCHL1, BD Biosciences), CD45RA-FITC (clone HI100, BioLegend), TCRb-BUV395 (clone IP26, BD Biosciences), HLA-DR-APC (clone Tu39, BioLegend)), PD-1-PECy7 (clone EH12-2H7, BioLegend), Granzyme B-PE (clone QA18A28, BioLegend). **Acquisition**: All the acquisitions were done on CytoFLEX LX (Beckman Coulter). FlowJo v10 was used for cytometry analysis. Percentages, geometric mean fluorescence intensities (gMFI), and absolute numbers (normalized to organ cellularity) were reported.

### Intravital cellular dynamics with 2-photon microscopy in the BM

For two-photon imaging experiments, LckGFP^hCD3ε^ chimeras were generated; for each timepoint, one to three mice were imaged, with 3-5 positions per mouse. Selected timepoints were D0 (baseline for 60 minutes before injection of CD20-TCE, and up to 240 minutes post-injection), D1, D2, D3 and D7 of TCE injection. At each timepoint, intravital imaging of the BM was realized as previously described^31^. Briefly, mice were premedicated with subcutaneous buprenorphin (Vetergesic, 0,1mg/kg) and were anesthetized with an intraperitoneal injection of ketamine (Imalgene 100mg/kg) + xylazine (Rompun, 10mg/kg). The scalp was removed, the skin was incised to expose the calvaria, and a coverslip was fixed above the frontoparietal suture. During imaging, mice were supplied with oxygen, and temperature was maintained at 37°C using a heated pad. **Recall experiments**: TCE was reinjected during imaging at prespecified timepoints (e.g., days 1, 3, 7); immediate changes in contact dynamics and apoptosis were quantified over the following 180 min. **Acquisition:** Two-photon imaging was performed with an upright microscope FVMPE-RS (OLYMPUS) and a 25X/1.05 NA water-dipping objective (OLYMPUS). Excitation was provided by an Insight DeepSee dual laser (Spectra Physics) tuned at 880 nm, typically used at 20% of laser power. The signal was first split through a 526nm long pass mirror; wavelengths < 526nm were split with a longpass 506 mirror, before filtration with filters 512/25 (for GFP) and 483/32 (for CFP) and detection on GaASP detectors; wavelengths > 526nm were split with a longpass 580 mirror, before filtration with filters 542/27 (for YFP) and 593/40 (for mTomato) and detection on standard detectors. To create time-lapse sequences, we typically scanned a 24-μm-thick volume of tissue at 3-μm Z-steps and 60-s intervals. **Image analysis:** Videos were processed and analyzed using the Fiji software (2.16.0/1.54p version; ImageJ2^71^). When necessary, images were stabilized using the Fast4Dreg^72,73^ Plugin. Tumor surface per frame was calculated using the Fiji software via AnalyzeParticles, after binarization of the image in the YFP channel using the Yen threshold. Characteristics of T cell movements, interactions and occurrence of mitosis, and tumor apoptosis, were quantified and scored manually. Contacts were defined *a priori* as the colocalization of T cell & tumor cell lasting for more than 3 minutes. They were annotated manually considering the duration (under vs over 10 minutes) and morphological characteristics of T cells (absence of T cell deformation and/or membrane overlap with the tumor cells were not considered stable contacts). Contact duration was evaluated over periods of 30 minutes before [-30;0 minutes] or after injection of TCE [+ 210;240 minutes] at D0. Redistribution of T cells on the tumor cells was assessed visually and quantified as the the nearest neighbor distance between tumor and T cell centroids. Briefly, for each intravital two-photon time-lapse recording, z-stacks were converted to maximum-intensity projections at each time point using Fiji. The centroid coordinates of tumor cells and T cells were manually extracted with Fiji on the first and last frame of the projected movie. A custom Python script^74^ computed for every tumor cell (Tumor*i*) its distance to the closest T cell in the XY plane among all T cells (T*j*), by calculating its Euclidean nearest-neighbor distance (NND*i*) defined as : *NNDi*=min*j*√(*xTj*−*xTumori*)2+(*yTj*−*yTumori*)2 where x and y are the *μ*m-calibrated spatial coordinates of each cell centroid in the projection plane. NND values were computed independently for the first and last frames. Data from 3 positions were pooled, each dot representing one tumor cell. Tumor apoptosis was quantified manually after blinding of the experimental timepoint, via the change of color resulting from FRET loss. Contact lethality was assessed as the ratio between pooled apoptosis events and the total count of contacts during the evaluation period (4 hours at D0 and 4 hours at D1, pool for 3 positions, in 1-3 mice). Cell density (apoptotic tumors or T cells) was quantified as the count of events per 1000μm^2^ of tumor cells, to allow comparison across positions and timepoints. Immediate killing post-TCE administration (within 3 hours of the 1^st^ injection or of a reinjection) was assessed as the accumulation of new apoptotic events normalized by the count of T cells in the field. Videos are shown as two-dimensional maximum intensity projections of three-dimensional data.

### Reprogramming of CD8^+^ T cells by sc-RNAseq

For scRNA-seq sample preparation, spleens from WT^hCD3ε^ mice bearing hCD20^+^DEVD^+^ Pro-B tumors were harvested at days D0, D3, D7, and D10 following TCE injection (n = 3 mice per group per timepoint), and mechanical dissociation was performed to obtain single cell suspension. Red blood cells were removed by Ficoll-Paque density gradient separation (Corning, cat. no. 25-072-CV), and the resulting hematopoietic cells were resuspended at a concentration of 20 × 10⁶ cells/mL in PBS supplemented with 0.04% BSA, without EDTA. For each mouse, 2 × 10⁶ cells were labeled with a distinct hashtag antibody according to the manufacturer’s protocol, using TotalSeq™-A0302 anti-mouse Hashtag Antibodies (clones M1/42; 30-F11, BioLegend). After labeling, the three biological replicates corresponding to the same experimental timepoint were pooled into a single low protein binding microcentrifuge tube (Thermo Scientific). Cell viability ranged between 68% and 83%. For each pool, 30,000 cells were immediately processed through the 10x Genomics GEM-X Universal 3′ Gene Expression v4 with Feature Barcoding workflow, using a Chromium X instrument (10x Genomics) and following the manufacturer’s User Guide (CG000731 – Revision A). Library quality and fragment size distribution were assessed using a Bioanalyzer 2100 (Agilent). Library concentrations were determined using a Qubit 2.0 fluorometer (Thermo Fisher Scientific). Libraries were sequenced on a NextSeq 2000 (Illumina) platform according to 10x Genomics recommendations. **Single-cell analysis:** For scRNAseq analysis, reads were processed with the 10x pipeline Cellranger 9.0.0 and the count command. We used the GRCm39 genome and transcriptome annotations provided by 10x Genomics. We added two sequences to this assembly. First, we added a consensus sequence of two fluorescent proteins [CFP, YFP], and human CD20 for identification of tumor cells. We also added the sequence and annotations of the plasmid pMSCV-hCD3ε-T2A-hCD34. We analyzed the data using R^75^ and the following packages: Seurat 5.5.0^76^, scDblFinder 1.26.0^77^, SingleR 2.14.0^78^ and monocle3 1.4.26^79^. From the raw Cellranger output, we selected cells with 2000 to 20823 (95th percentile of the distribution) unique molecular identifiers (UMIs), more than 1000 detected feature, less than 20% of mitochondrial UMIs. We removed the cells containing the hashtag corresponding to more than one replicate and cells detected as doublets (D0: 9251 cells, D3: 8559, D7: 7972, D10: 9425). We then focused on the time point D0, D3 and D7. We defined the S and G2M scores from the mouse orthologs of the “S phase” genes and “G2M” genes provided in the Seurat pipeline. We processed them in 3 steps: 1) selection of T cells, 2) selection of CD8 and 3) clustering of the CD8^+^ T cells. In each step, we detected and removed doublet cells, scaled the data while considering the S and G2M scores. We also used 50 principal components (PCs) for neighbor finding, clustering and generating the Uniform Manifold Approximation Projection (UMAP). We selected the resolution parameter of the FindClusters function by checking the number of cells by cluster and the clustree. For the step 1 and 2 we annotated each cluster using the normalized expression values of 830 microarray samples of pure mouse immune cells^80^ and the SingleR package. For step 1 the resolution was 1.1 and we selected the cluster annotated “T cells”. For step 2 the resolution was 0.7, we removed any cluster not annotated “T cells”, kept the cluster with at least 50% of cells expressing *Cd8a* or *Cd8b1*, but removed the cells expressing ≥ 1 *Cd4* read. For step 3 the resolution was 0.7 and the resulting umap was used to compute trajectories in the package Monocle3. To find the genes representing each CD8 T cells cluster, we used the Wilcoxon Rank Sum test on the normalized data as implemented in the function FindAllMarkers in Seurat.

### Remodeling of the chemokine landscape by multiplex assays

Bone marrow homogenates from flushed tibia & fibula and spleen homogenates were collected at day 1 and day 3 post-TCE or isotype control. Cells were resuspended in 40 µL/10^6^ of cells of Lysis Buffer (Invitrogen) containing Protease Cocktail Inhibitor (1:100, Thermo Fisher) and Pierce Universal Nuclease. Then, cell lysates were briefly sonicated (30 seconds at 25 KHz). Then, the supernatant was collected by ultracentrifugation at 14,000 g for 10 minutes in a cold microfuge. Multiplex assays were conducted using the 20-Plex ProcartPlex Panel (Mouse Th1/Th2 Cytokine & Chemokine 20plex, from Invitrogen) according to the manufacturer’s instructions. Data analysis was conducted using a Bio-Plex 200 system with Bio-Plex Manager software (Bio-Rad). Standard curves and QC samples ensured assay performance.

### Exploration of the generation of CTL_TCE_ via adoptive transfer

T cell subsets were sorted from spleen and LNs of LckGFP^hCD3ε^ mice, using 2 successive sorting methods: 1/column enrichment of hCD34^+^ T cells using negative T cell (Pan T Cells Isolation Kit II Mouse cat #130-095-130) and positive hCD34 (MicroBeads Kit cat #130-046-702, from Miltenyi) selections. 2/ The enriched hCD34+ T cells were then FACS sorted on an BD FACS ARIA III based on CD62L and CD44 expression. Specifically, we sorted T_Naïve_ (CD62L^+^ CD44^-^), T_CM_ (CD62L^+^ CD44^+^) and T_EM_ (CD62L^-^ CD44^+^) populations. A purity > 95% was obtained. Equal counts of sorted cell subsets were then transferred into non-chimeric hCD20^+^DEVD^+^ Pro-B-bearing C57BL/6 recipients (5×10^5^ cells per mouse). Six groups of 4 mice were constituted in total (pooled from 2 independent experiments), where recipient mice received either T_Naïve_, T_CM_ or T_EM_ subsets, before treatment on the following day either with TCE or isotype control. Six days after adoptive transfer, spleen, blood, and lymph nodes were harvested, and hCD3ε GFP^+^ T cells were recovered and analyzed by flow cytometry.

### Survival experiments

WT^hCD3ε^ or Xcr1-DTA^hCD3ε^ mice were injected with tumor cells, hCD20^+^DEVD^+^ Pro-B (or hCD20^+^DEVD^+^OVA^+^ Pro-B for evaluation of cDC1 role in long-term tumor control). Three days after tumor cell injection, mice were treated with one injection of CD20-TCE or isotype control as described above. For rechallenge experiments, WT^hCD3ε^ mice that had achieved complete clinical remission after TCE were rechallenged with hCD20^+^DEVD^+^ Pro-B cells, without additional therapy; age-matched controls were included. Mouse behavior was evaluated three times per week, and mice were euthanized upon reaching humane endpoints: general appearance, body condition score (BCS), locomotion and behavior, respiratory distress (assessed by in-cage observation).

### Statistical analysis

All statistical analyses were conducted using Prism V 11.0.2 (GraphPad). Data are presented as Mean ± SEM. Two-tailed unpaired t test with Welch’s correction was used for comparisons between two groups, while one-way ANOVA with Tukey’s adjustment for multiple comparison. For mice survival, statistical analysis was performed using a log-rank test, with Holm-Šídák’s multiple comparisons test when required. All statistical tests were two-tailed with a significance level of 0.05; ns, non-significant.

## Data availability

Single-cell RNA-sequencing data and code availability will be deposited upon publication. Other data are available from the corresponding author upon reasonable request.

## Acknowledgments

We are deeply grateful to Philippe Bousso, who conceived and supervised this work until his passing. His vision of *in vivo* immunology shaped every aspect of this study.

We thank members of the Bousso laboratory for critical review of the manuscript, Nicolas Serafini and Sébastien Lemoine for helpful discussions. We thank Bérengère Hugot for her administrative support. We are grateful for the mice that were used to perform these experiments, and for the human donors from whom we obtained PBMCs. We acknowledge the mouse facility and CB UTechS at Institut Pasteur for support in conducting the present study. We thank the Biomics Platform, C2RT, Institut Pasteur, Paris, France, and more specifically Florence Jagorel, supported by France Génomique (ANR-10-INBS-09) and IBISA, for assistance in single-cell RNA-sequencing experiments. We thank Genentech for providing the CD20-TCE molecule under a material transfer agreement. We thank the NIH Tetramer Core Facility for providing H2-K^b^-OVA tetramers. We acknowledge BioRender from which icons were sourced to generate experimental setup schematics. The work was supported by Institut Pasteur, INSERM, La Fondation ARC (project ARPGA2022120005636_6359), and an Advanced grant (ENLIGHTEN) from the European Research Council (PB). MML was supported by an “Année recherche” fellowship from Assistance Publique-Hôpitaux de Paris & Agence Régionale de Santé, and two doctoral grants from Fondation pour la Recherche Médicale “FDM-Poste de thèse pour internes et assistants” and “ARC_Fin de thèse ARC_FRM”.

## Author contributions

PB conceptualized the project and secured funding. MML & PB designed the experiments. MML & BC conducted the experiments with ZG (2-photon intravital imaging), RA (FACS sorting), CMC & VS (scRNAseq). FL generated the cell lines and plasmids and managed orders for reagents and consumables. LCF contributed to daily organization of lab business. CG wrote the ethical project for authorization of experimental procedures and ensured respect for animal welfare.

MML & PB analyzed the data with support from AV & CG (single-cell RNA-seq analysis) and BA (NND, Distance analysis of intravital imaging). CG, MVG & PG provided important scientific insight. MML & PB wrote the manuscript.

## References

1. Baeuerle, P. A. & Reinhardt, C. Bispecific T-Cell Engaging Antibodies for Cancer Therapy: Figure 1. Cancer Res. 69, 4941–4944 (2009).

2. Goebeler, M.-E. & Bargou, R. C. T cell-engaging therapies — BiTEs and beyond. Nat. Rev. Clin. Oncol. 17, 418–434 (2020).

3. Slaney, C. Y., Wang, P., Darcy, P. K. & Kershaw, M. H. CARs versus BiTEs: A Comparison between T Cell–Redirection Strategies for Cancer Treatment. Cancer Discov. 8, 924–934 (2018).

4. Labrijn, A. F., Janmaat, M. L., Reichert, J. M. & Parren, P. W. H. I. Bispecific antibodies: a mechanistic review of the pipeline. Nat. Rev. Drug Discov. 18, 585–608 (2019).

5. Cappell, K. M. & Kochenderfer, J. N. Long-term outcomes following CAR T cell therapy: what we know so far. Nat. Rev. Clin. Oncol. 20, 359–371 (2023).

6. Neelapu, S. S. et al. Axicabtagene Ciloleucel CAR T-Cell Therapy in Refractory Large B-Cell Lymphoma. N. Engl. J. Med. 377, 2531–2544 (2017).

7. Bargou, R. et al. Tumor Regression in Cancer Patients by Very Low Doses of a T Cell–Engaging Antibody. Science 321, 974–977 (2008).

8. Sun, L. L. et al. Anti-CD20/CD3 T cell–dependent bispecific antibody for the treatment of B cell malignancies. Sci. Transl. Med. 7, (2015).

9. Budde, L. E. et al. Safety and efficacy of mosunetuzumab, a bispecific antibody, in patients with relapsed or refractory follicular lymphoma: a single-arm, multicentre, phase 2 study. Lancet Oncol. 23, 1055–1065 (2022).

10. Dickinson, M. J. et al. Glofitamab for Relapsed or Refractory Diffuse Large B-Cell Lymphoma. N. Engl. J. Med. 387, 2220–2231 (2022).

11. Kantarjian, H. et al. Blinatumomab versus Chemotherapy for Advanced Acute Lymphoblastic Leukemia. N. Engl. J. Med. 376, 836–847 (2017).

12. Moreau, P. et al. Teclistamab in Relapsed or Refractory Multiple Myeloma. N. Engl. J. Med. 387, 495–505 (2022).

13. Thieblemont, C. et al. Epcoritamab in relapsed/refractory large B-cell lymphoma: 2-year follow-up from the pivotal EPCORE NHL-1 trial. Leukemia 38, 2653–2662 (2024).

14. Schmeing, S. et al. Molecular Features of Response and Resistance to Glofitamab, a T-Cell Engager for treatment of Large B-Cell Lymphoma. Blood Adv. bloodadvances.2025016925 (2026) doi:10.1182/bloodadvances.2025016925.

15. Zannikou, M. et al. Bi-specific T cell-engaging antibody triggers protective immune memory and glioma microenvironment remodeling in immune-competent preclinical models. J. Immunother. Cancer 13, e011714 (2025).

16. Eissenberg, L. G. et al. Control of acute myeloid leukemia and generation of immune memory in vivo using AMV564, a bivalent bispecific CD33 x CD3 T cell engager. PLOS ONE 19, e0300174 (2024).

17. Kroemer, G., Galluzzi, L., Kepp, O. & Zitvogel, L. Immunogenic Cell Death in Cancer Therapy. Annu. Rev. Immunol. 31, 51–72 (2013).

18. Dreier, T. et al. Extremely potent, rapid and costimulation-independent cytotoxic T-cell response against lymphoma cells catalyzed by a single-chain bispecific antibody. Int. J. Cancer 100, 690–697 (2002).

19. Ellerman, D. Bispecific T-cell engagers: Towards understanding variables influencing the in vitro potency and tumor selectivity and their modulation to enhance their efficacy and safety. Methods 154, 102–117 (2019).

20. Gruen, M., Bommert, K. & Bargou, R. C. T-cell-mediated lysis of B cells induced by a CD19xCD3 bispecific single-chain antibody is perforin dependent and death receptor independent. Cancer Immunol. Immunother. 53, 625–632 (2004).

21. Hoffmann, P. et al. Serial killing of tumor cells by cytotoxic T cells redirected with a CD19-/CD3-bispecific single-chain antibody construct. Int. J. Cancer 115, 98–104 (2005).

22. Offner, S., Hofmeister, R., Romaniuk, A., Kufer, P. & Baeuerle, P. A. Induction of regular cytolytic T cell synapses by bispecific single-chain antibody constructs on MHC class I-negative tumor cells. Mol. Immunol. 43, 763–771 (2006).

23. Corrado, F. et al. Single-Cell Transcriptomic Profiling of T Cells from Blinatumomab-Treated Patients with B-Cell Precursor Acute Lymphoblastic Leukemia (BCP-ALL) Reveals Circulating CD8 T Cell Subsets Associated with Treatment Response. Blood 142, 600–600 (2023).

24. Friedrich, M. J. et al. The pre-existing T cell landscape determines the response to bispecific T cell engagers in multiple myeloma patients. Cancer Cell S1535610823000363 (2023) doi:10.1016/j.ccell.2023.02.008.

25. Huo, Y. et al. Blinatumomab-induced T cell activation at single cell transcriptome resolution. BMC Genomics 22, 145 (2021).

26. Falchi, L. et al. Intratumoral T-cell composition predicts epcoritamab-based treatment efficacy in B-cell non-Hodgkin lymphomas. Preprint at 10.1101/2024.07.02.24309792 (2024).

27. Cremasco, F. et al. Cross-linking of T cell to B cell lymphoma by the T cell bispecific antibody CD20-TCB induces IFNγ/CXCL10-dependent peripheral T cell recruitment in humanized murine model. PLOS ONE 16, e0241091 (2021).

28. Ueda, O. et al. Entire CD3ε, δ, and γ humanized mouse to evaluate human CD3–mediated therapeutics. Sci. Rep. 7, 45839 (2017).

29. Zorn, J. A. et al. Humanization of a strategic CD3 epitope enables evaluation of clinical T-cell engagers in a fully immunocompetent in vivo model. Sci. Rep. 12, 3530 (2022).

30. de la Hera, A., Müller, U., Olsson, C., Isaaz, S. & Tunnacliffe, A. Structure of the T cell antigen receptor (TCR): two CD3 epsilon subunits in a functional TCR/CD3 complex. J. Exp. Med. 173, 7–17 (1991).

31. Breart, B., Lemaître, F., Celli, S. & Bousso, P. Two-photon imaging of intratumoral CD8+ T cell cytotoxic activity during adoptive T cell therapy in mice. J. Clin. Invest. 118, 1390–1397 (2008).

32. Galeano Niño, J. L., et al. Cytotoxic T cells swarm by homotypic chemokine signalling. eLife 9, e56554 (2020).

33. Liao, C.-Y. et al. CD3-engaging bispecific antibodies trigger a paracrine regulated wave of T-cell recruitment for effective tumor killing. *Commun*. Biol. 7, 983 (2024).

34. Sallusto, F., Geginat, J. & Lanzavecchia, A. Central Memory and Effector Memory T Cell Subsets: Function, Generation, and Maintenance. Annu. Rev. Immunol. 22, 745–763 (2004).

35. Stinchcombe, J. C. & Griffiths, G. M. Secretory Mechanisms in Cell-Mediated Cytotoxicity. Annu. Rev. Cell Dev. Biol. 23, 495–517 (2007).

36. Voskoboinik, I., Whisstock, J. C. & Trapani, J. A. Perforin and granzymes: function, dysfunction and human pathology. Nat. Rev. Immunol. 15, 388–400 (2015).

37. Ferrini, S. et al. The LFA-1/ICAM cell adhesion pathway is involved in tumor-cell lysis mediated by bispecific monoclonal-antibody-targeted T lymphocytes. Int. J. Cancer 56, 846–852 (1994).

38. Dustin, M. L. The Immunological Synapse. Cancer Immunol. Res. 2, 1023–1033 (2014).

39. Leithner, A., et al. Solution structure and synaptic analyses reveal determinants of bispecific T cell engager potency. Proc. Natl. Acad. Sci. 122, e2425781122 (2025).

40. Dominguez, C. X. et al. The transcription factors ZEB2 and T-bet cooperate to program cytotoxic T cell terminal differentiation in response to LCMV viral infection. J. Exp. Med. 212, 2041–2056 (2015).

41. Omilusik, K. D. et al. Transcriptional repressor ZEB2 promotes terminal differentiation of CD8+ effector and memory T cell populations during infection. J. Exp. Med. 212, 2027–2039 (2015).

42. Im, S. J. et al. Defining CD8+ T cells that provide the proliferative burst after PD-1 therapy. Nature 537, 417–421 (2016).

43. Miller, B. C. et al. Subsets of exhausted CD8+ T cells differentially mediate tumor control and respond to checkpoint blockade. Nat. Immunol. 20, 326–336 (2019).

44. Siddiqui, I. et al. Intratumoral Tcf1+PD-1+CD8+ T Cells with Stem-like Properties Promote Tumor Control in Response to Vaccination and Checkpoint Blockade Immunotherapy. Immunity 50, 195–211.e10 (2019).

45. Beltra, J.-C. et al. Developmental Relationships of Four Exhausted CD8+ T Cell Subsets Reveals Underlying Transcriptional and Epigenetic Landscape Control Mechanisms. Immunity 52, 825–841.e8 (2020).

46. Gattinoni, L. et al. A human memory T cell subset with stem cell–like properties. Nat. Med. 17, 1290–1297 (2011).

47. Lugli, E. et al. Superior T memory stem cell persistence supports long-lived T cell memory. J. Clin. Invest. JCI66327 (2013) doi:10.1172/JCI66327.

48. Fraietta, J. A. et al. Determinants of response and resistance to CD19 chimeric antigen receptor (CAR) T cell therapy of chronic lymphocytic leukemia. Nat. Med. 24, 563–571 (2018).

49. Hiraga, J. et al. Down-regulation of CD20 expression in B-cell lymphoma cells after treatment with rituximab-containing combination chemotherapies: its prevalence and clinical significance. Blood 113, 4885–4893 (2009).

50. Betts, A. & Van Der Graaf, P. H. Mechanistic Quantitative Pharmacology Strategies for the Early Clinical Development of Bispecific Antibodies in Oncology. Clin. Pharmacol. Ther. 108, 528–541 (2020).

51. Zieger, N. et al. T-cell exhaustion induced by continuous bispecific molecule exposure is ameliorated by treatment-free intervals. Blood blood.2022015956 (2022) doi:10.1182/blood.2022015956.

52. Macián, F. et al. Transcriptional Mechanisms Underlying Lymphocyte Tolerance. Cell 109, 719–731 (2002).

53. Martinez, G. J. et al. The Transcription Factor NFAT Promotes Exhaustion of Activated CD8 + T Cells. Immunity 42, 265–278 (2015).

54. Scott, A. C. et al. TOX is a critical regulator of tumour-specific T cell differentiation. Nature 571, 270–274 (2019).

55. Khan, O. et al. TOX transcriptionally and epigenetically programs CD8+ T cell exhaustion. Nature 571, 211–218 (2019).

56. Ma, H. et al. Combination therapy with T cell engager and PD-L1 blockade enhances the antitumor potency of T cells as predicted by a QSP model. J. Immunother. Cancer 8, e001141 (2020).

57. Chen, D. S. & Mellman, I. Oncology Meets Immunology: The Cancer-Immunity Cycle. Immunity 39, 1–10 (2013).

58. Garris, C. S. et al. Successful Anti-PD-1 Cancer Immunotherapy Requires T Cell-Dendritic Cell Crosstalk Involving the Cytokines IFN-γ and IL-12. Immunity 49, 1148–1161.e7 (2018).

59. Galluzzi, L. et al. Consensus guidelines for the definition, detection and interpretation of immunogenic cell death. J. Immunother. Cancer 8, e000337 (2020).

60. Budde, L. E. et al. Single-Agent Mosunetuzumab Shows Durable Complete Responses in Patients With Relapsed or Refractory B-Cell Lymphomas: Phase I Dose-Escalation Study. J. Clin. Oncol. JCO.21.00931 (2021) doi:10.1200/JCO.21.00931.

61. Zitvogel, L., Apetoh, L., Ghiringhelli, F. & Kroemer, G. Immunological aspects of cancer chemotherapy. Nat. Rev. Immunol. 8, 59–73 (2008).

62. Böttcher, J. P. & Reis E Sousa, C. The Role of Type 1 Conventional Dendritic Cells in Cancer Immunity. Trends Cancer 4, 784–792 (2018).

63. Hildner, K., et al. *Batf3* Deficiency Reveals a Critical Role for CD8α^+^ Dendritic Cells in Cytotoxic T Cell Immunity. Science 322, 1097–1100 (2008).

64. Roberts, E. W. et al. Critical Role for CD103+/CD141+ Dendritic Cells Bearing CCR7 for Tumor Antigen Trafficking and Priming of T Cell Immunity in Melanoma. Cancer Cell 30, 324–336 (2016).

65. Salmon, H. et al. Expansion and Activation of CD103+ Dendritic Cell Progenitors at the Tumor Site Enhances Tumor Responses to Therapeutic PD-L1 and BRAF Inhibition. Immunity 44, 924–938 (2016).

66. Middelburg, J. et al. T-cell stimulating vaccines empower CD3 bispecific antibody therapy in solid tumors. Nat. Commun. 15, 48 (2024).

67. Poussin, M. et al. Dichotomous impact of affinity on the function of T cell engaging bispecific antibodies. J. Immunother. Cancer 9, e002444 (2021).

68. Middelburg, J. et al. Overcoming Challenges for CD3-Bispecific Antibody Therapy in Solid Tumors. Cancers 13, 287 (2021).

69. You, R. et al. Visualizing Spatial and Stoichiometric Barriers to Bispecific T-cell Engager Efficacy. Cancer Immunol. Res. canimm.CIR-21–0594-E.2021 (2022) doi:10.1158/2326-6066.CIR-21-0594.

70. Naviaux, R. K., Costanzi, E., Haas, M. & Verma, I. M. The pCL vector system: rapid production of helper-free, high-titer, recombinant retroviruses. J. Virol. 70, 5701–5705 (1996).

71. Rueden, C. T. et al. ImageJ2: ImageJ for the next generation of scientific image data. BMC Bioinformatics 18, 529 (2017).

72. Pylvänäinen, J. W. et al. Fast4DReg – fast registration of 4D microscopy datasets. J. Cell Sci. 136, jcs260728 (2023).

73. Laine, R. F. et al. NanoJ: a high-performance open-source super-resolution microscopy toolbox. J. Phys. Appl. Phys. 52, 163001 (2019).

74. BptisteArnaud. BptisteArnaud/nnd-analysis: NND Analysis v1.0.1 - Zenodo archive sync. Zenodo 10.5281/ZENODO.21256807 (2026).

75. R Core Team. R: A Language and Environment for Statistical Computing. R Foundation for Statistical Computing (2025).

76. Hao, Y. et al. Dictionary learning for integrative, multimodal and scalable single-cell analysis. Nat. Biotechnol. 42, 293–304 (2024).

77. Germain, P.-L., Lun, A., Garcia Meixide, C., Macnair, W. & Robinson, M. D. Doublet identification in single-cell sequencing data using scDblFinder. F1000Research 10, 979 (2021).

78. Aran, D. et al. Reference-based analysis of lung single-cell sequencing reveals a transitional profibrotic macrophage. Nat. Immunol. 20, 163–172 (2019).

79. Qiu, X. et al. Reversed graph embedding resolves complex single-cell trajectories. Nat. Methods 14, 979–982 (2017).

80. Heng, T. S. P., Painter, M. W., & Immunological Genome Project Consortium. The Immunological Genome Project: networks of gene expression in immune cells. Nat. Immunol. 9, 1091–1094 (2008).

