## Extended data for "T cell engagers convert acute tumor killing into endogenous vaccination"

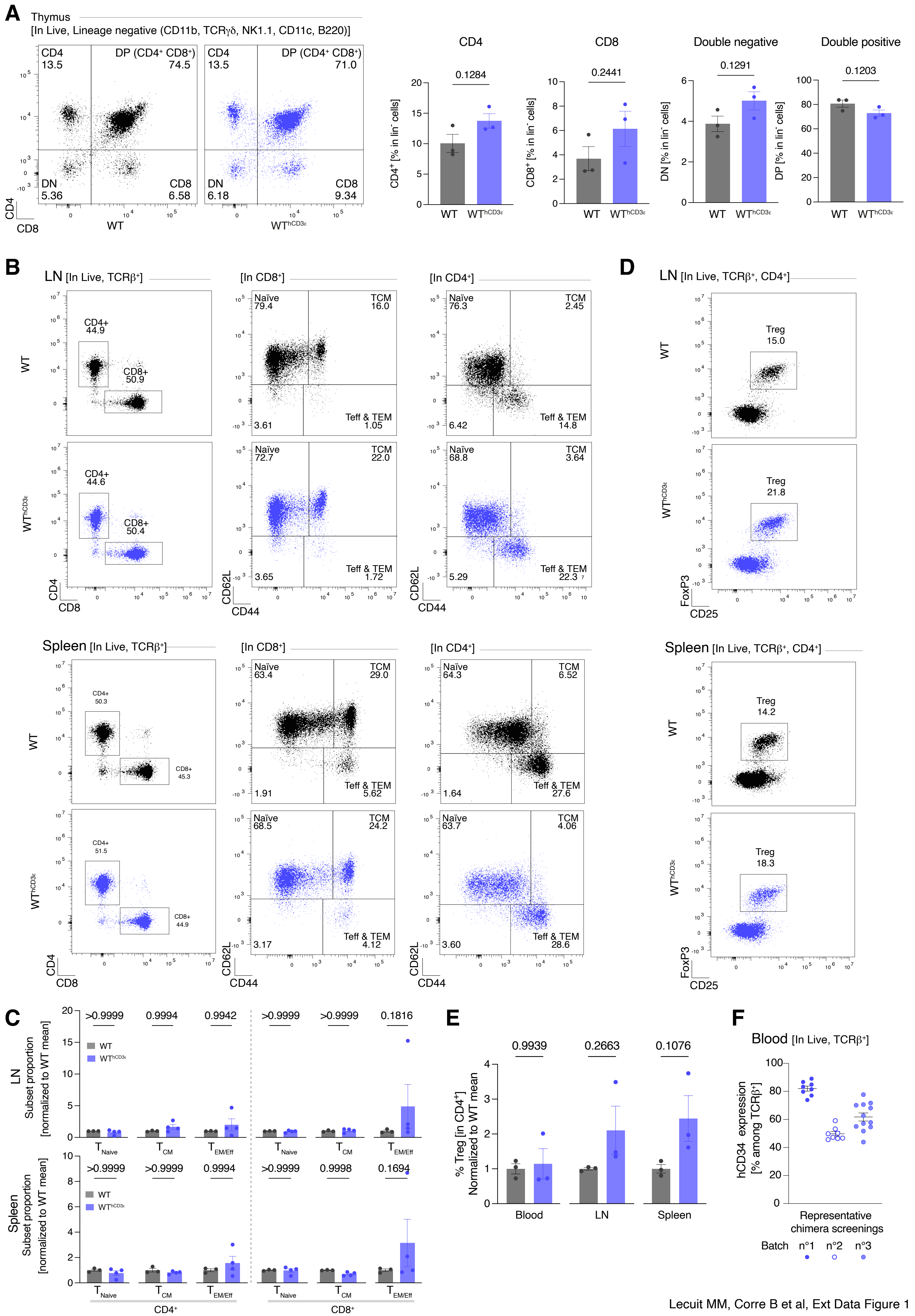


Lecuit MM, Corre B et al, Ext Data Figure 1

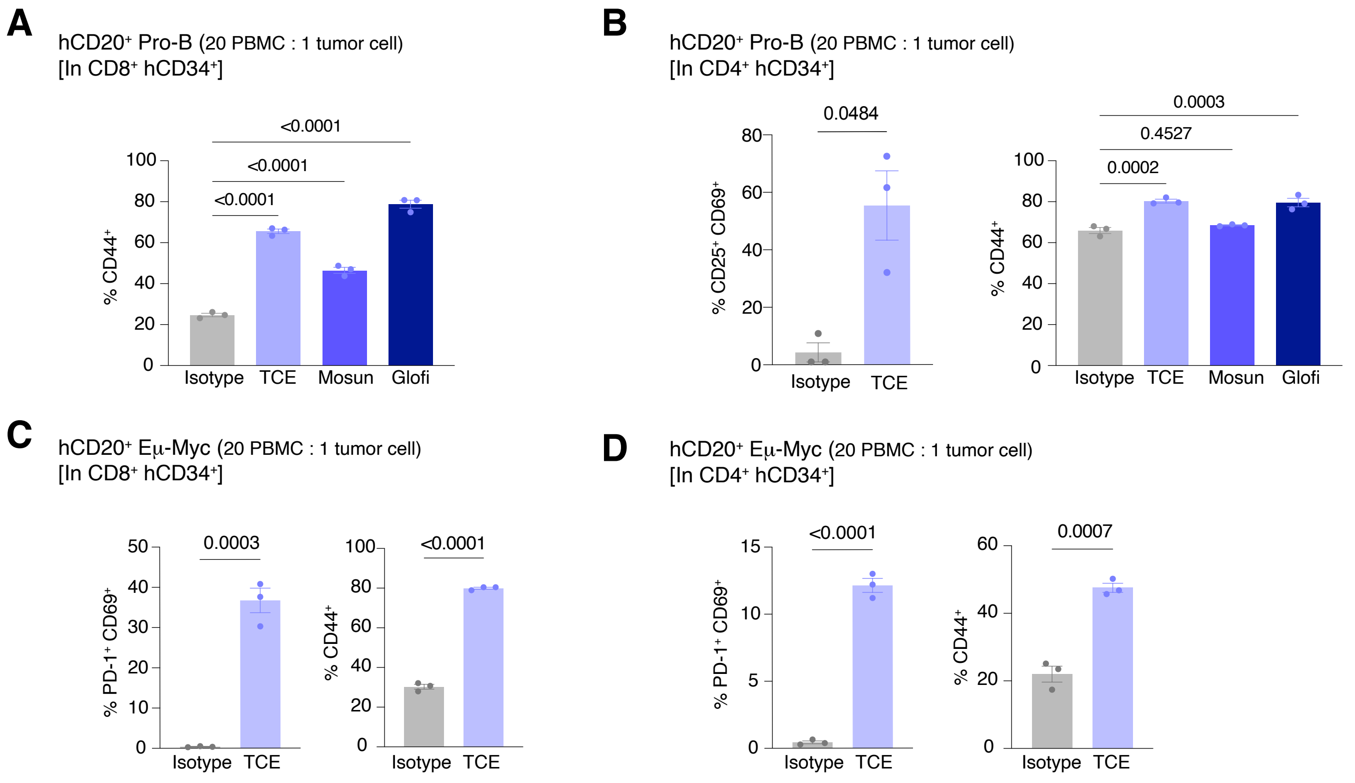


Lecuit MM, Corre B et al, Ext Data Figure 2

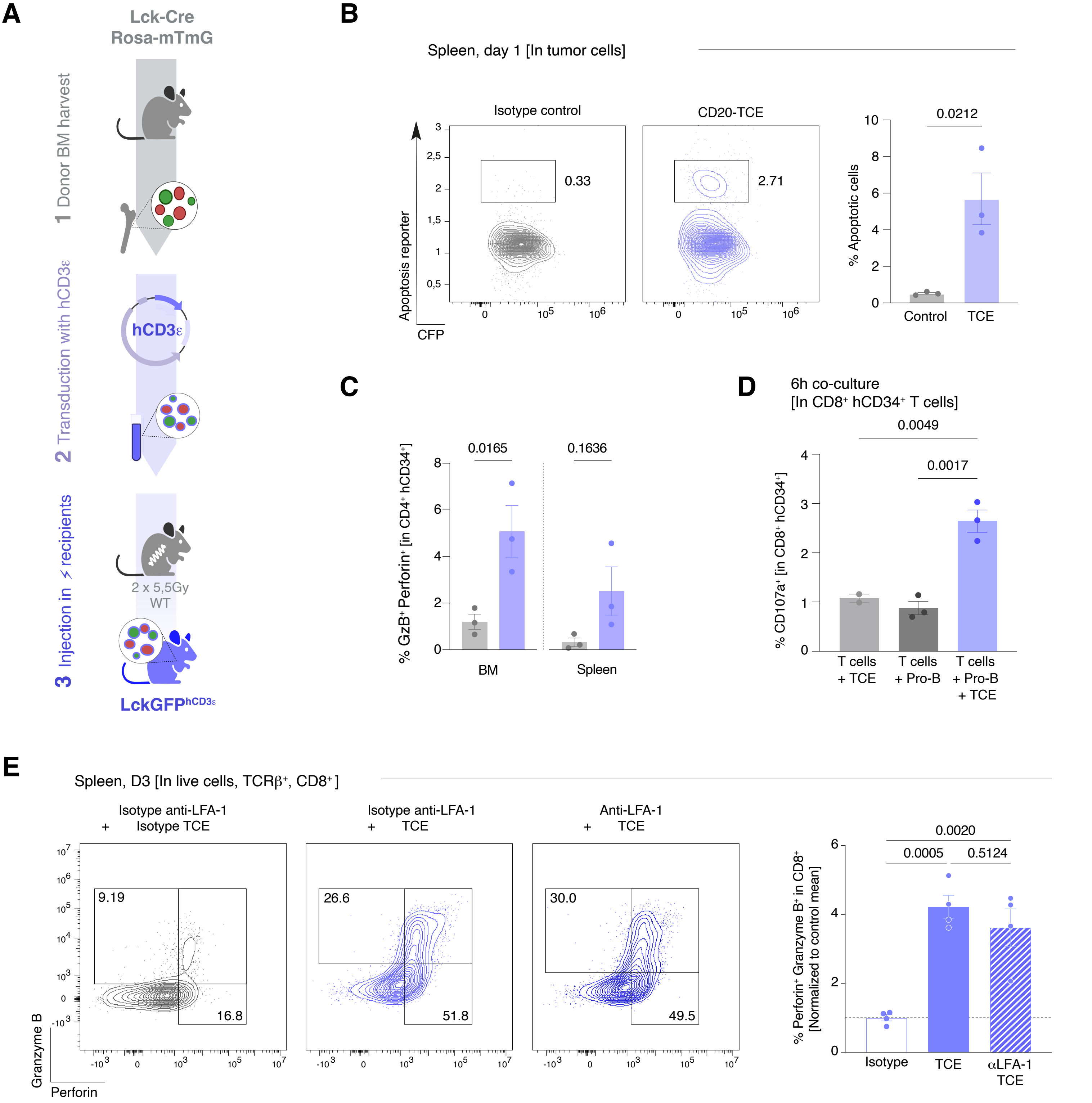


Lecuit MM, Corre B et al, Ext Data Figure 3

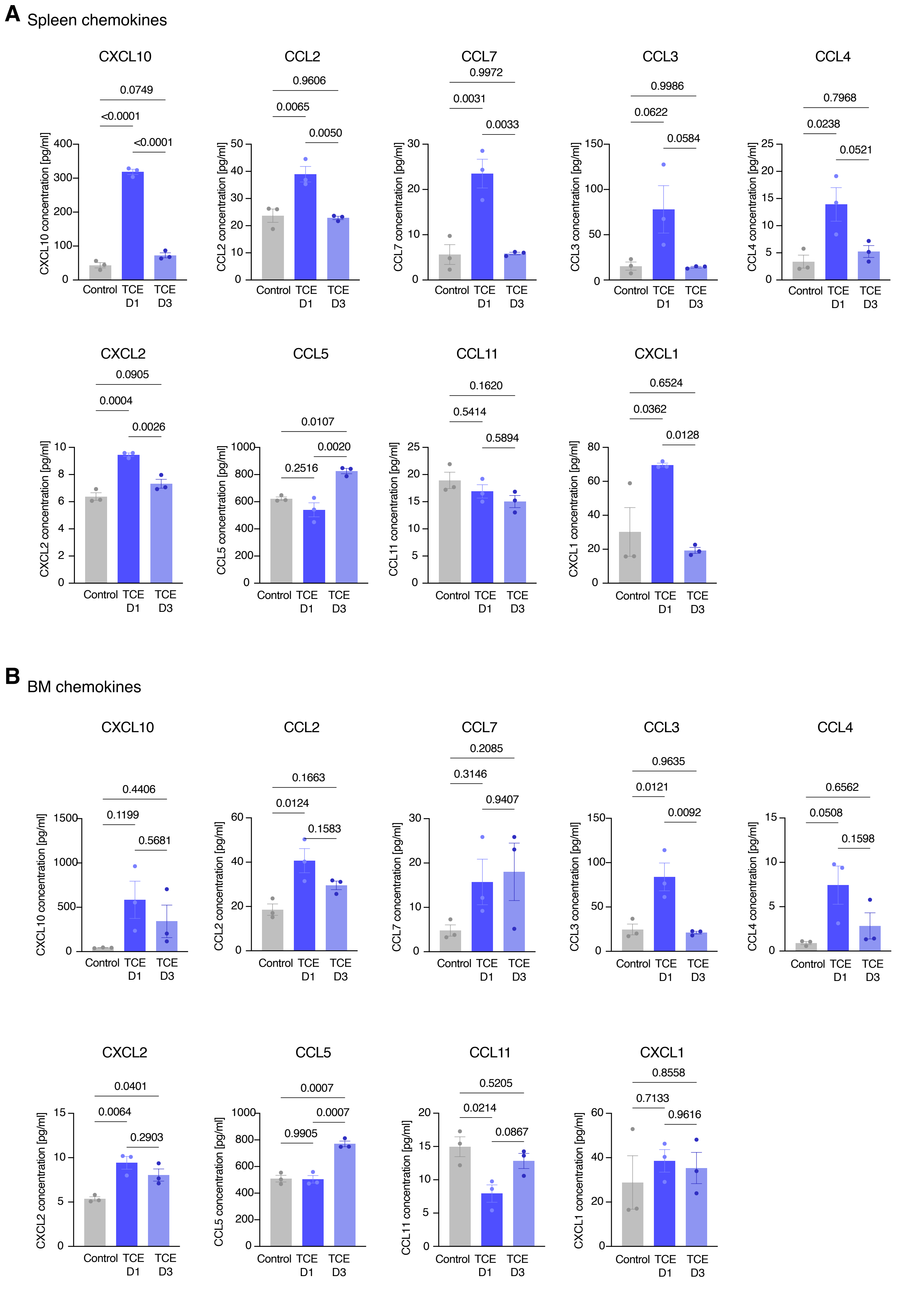


Lecuit MM, Corre B et al, Ext Data Figure 4

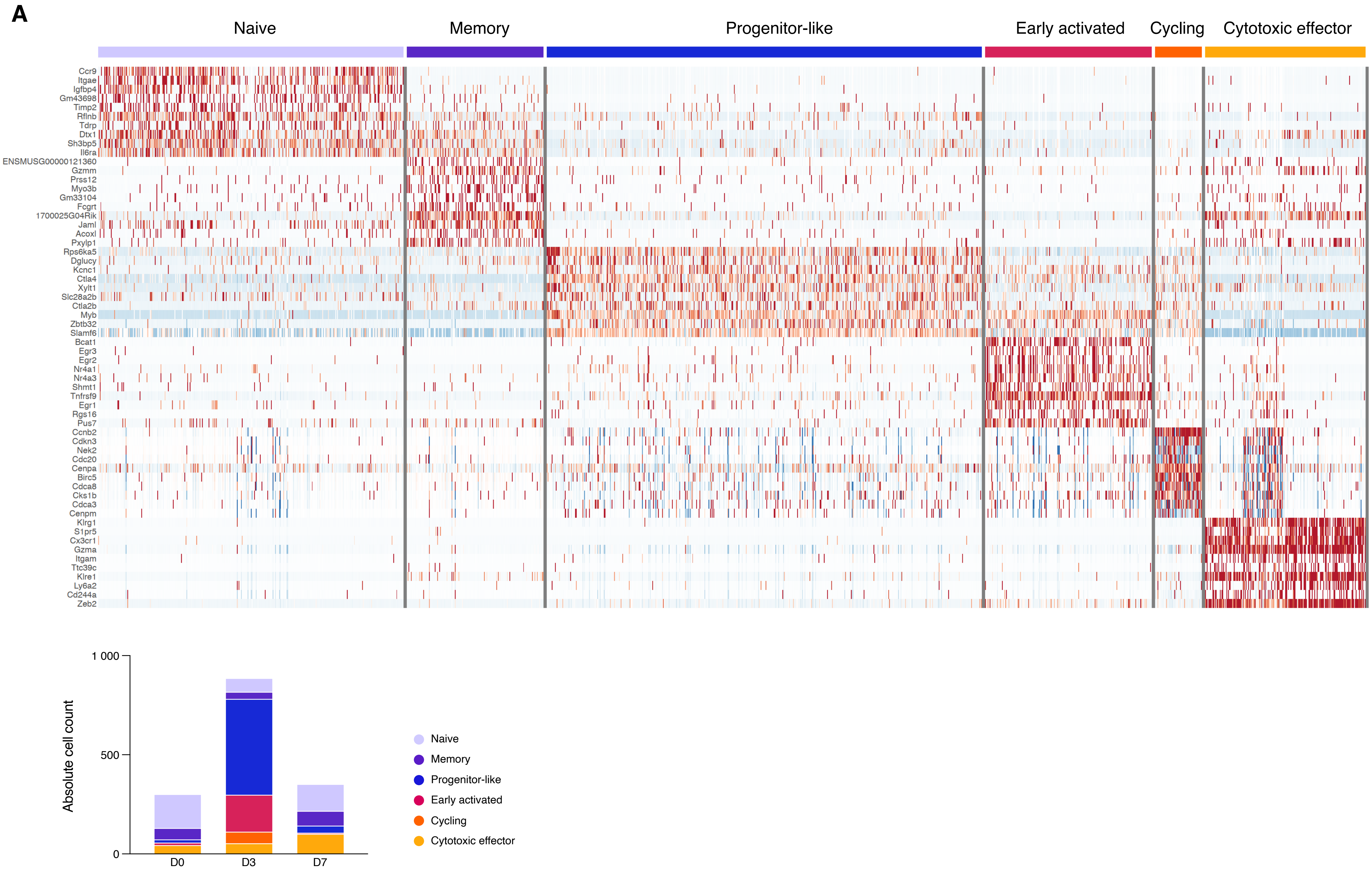


Lecuit MM, Corre B et al, Ext Data Figure 5

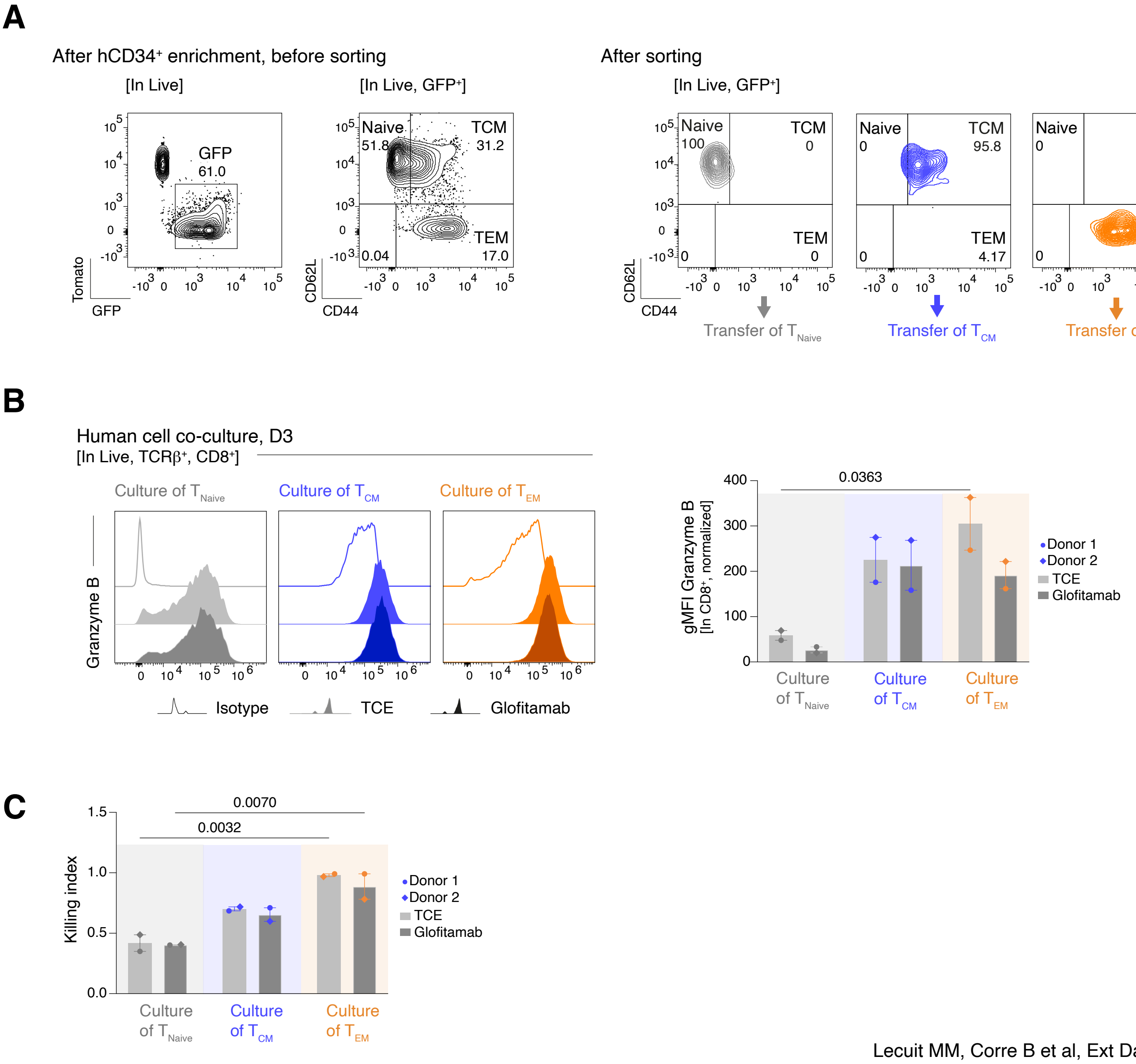


Lecuit MM, Corre B et al, Ext Data Figure 6

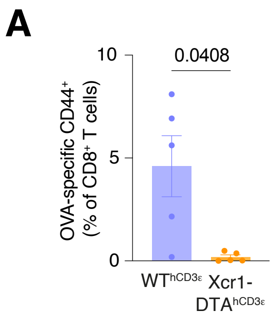


Lecuit MM, Corre B et al, Ext Data Figure 7
